# Plasmacytoid dendritic cells regulate megakaryocyte and platelet homeostasis

**DOI:** 10.1101/2022.05.31.494147

**Authors:** Florian Gaertner, Hellen Ishikawa-Ankerhold, Susanne Stutte, Wenwen Fu, Chenglong Guo, Jutta Weitz, Anne Dueck, Zhe Zhang, Dominic van den Heuvel, Valeria Fumagalli, Michael Lorenz, Louisa von Baumgarten, Konstantin Stark, Tobias Straub, Saskia von Stillfried, Peter Boor, Marco Colonna, Christian Schulz, Thomas Brocker, Barbara Walzog, Christoph Scheiermann, Stefan Engelhardt, William C. Aird, Tobias Petzold, Michael Sixt, Martina Rudelius, Claus Nerlov, Matteo Iannacone, Robert A. J. Oostendorp, Steffen Massberg

## Abstract

Platelet homeostasis is essential for vascular integrity and immune defense. While the process of platelet formation by fragmenting megakaryocytes (thrombopoiesis) has been extensively studied, the cellular and molecular mechanisms required to constantly replenish the pool of megakaryocytes by their progenitor cells (megakaryopoiesis) remains unclear. Here we use intravital 2 photon microscopy to track individual megakaryopoiesis over days. We identify plasmacytoid dendritic cells (pDCs) as crucial bone marrow niche cells that regulate megakaryopoiesis. pDCs monitor the bone marrow for platelet-producing megakaryocytes and deliver IFN-α to the megakaryocytic niche to trigger local on-demand proliferation of megakaryocyte progenitors. This fine-tuned coordination between thrombopoiesis and megakaryopoiesis is crucial for megakaryocyte and platelet homeostasis in steady state and stress. However, uncontrolled pDC function within the megakaryocytic niche is detrimental. Accordingly, we show that pDCs activated by SARS-CoV2 drive inappropriate megakaryopoiesis associated with thrombotic complications. Together, we uncover a hitherto unknown megakaryocytic bone marrow niche maintained by the constitutive delivery of pDC-derived IFN-α.

## Main

The hematopoietic system comprises a heterogeneous population of terminally differentiated cells that fulfill distinct roles in host protection and tissue oxygenation. Platelets are terminally differentiated hematopoietic cells that form clots and prevent blood loss after vascular injury^1^. They also assist immune cells and fight pathogen intruders in a process referred to as immunothrombosis^2^. Platelets are produced in the bone marrow (BM) by their precursors, the megakaryocytes (MKs) in a process termed thrombopoiesis^3^. During thrombopoiesis, MKs release fragments while their cell body is entirely consumed^4^. Consequently, replenishment of fragmented MKs from their MK progenitors (termed megakaryopoiesis) is required to ensure sufficient platelet production.

Here, we used two-photon intravital microscopy (2P-IVM) of the megakaryocytic lineage to identify the spatio-temporal patterns of megakaryopoiesis during steady state and in response to thrombocytopenia. We show that thrombopoiesis is associated with MK consumption, a process that is compensated by continuous local proliferation of MK progenitors within the megakaryocytic niche. We establish that BM plasmacytoid dendritic cells (pDCs), a unique subset of dendritic innate immune cells specialized in antiviral immunity^5^, constitute crucial niche cells controlling MK homeostasis. pDCs constantly scan BM tissue and secrete Interferon alpha (IFN-α) upon detection of exhausted MKs. We demonstrate that local pDC-derived IFN-α synchronizes megakaryopoiesis and thrombopoiesis to maintain homeostasis of MKs and platelets in steady state and also accounts for enhanced platelet consumption in disease. Viral infection with SARS-CoV-2 can manipulate pDC-driven MK proliferation leading to inappropriate megakaryopoiesis, which has been associated with thrombotic complications during COVID-19^6^.

### The megakaryocytic BM niche spatio-temporally coordinates megakaryopoiesis and thrombopoiesis

The primary site of megakaryopoiesis in mammals is the BM^3^. To analyze the spatial distribution of MKs (CD41^+^; CD42^+^) and their progenitors (MKPs) (CD41^+^; CD42^-^) we performed three-dimensional immunofluorescence imaging of mouse BM^7^ (**Fig. 1a, Extended Fig. 1a and Supplementary Video 1**). The vast majority of mature MKs (∼82%) resides within a distance of 5 µm to sinusoids (**Extended Data Fig. 1b**) without preferential association to the endosteum (**Extended Data Fig. 1c**). MKPs were significantly smaller than MKs (**Extended Data Fig. 1d**) and showed a largely spherical shape (**Extended Data Fig. 1e**). They have a distribution similar to mature MKs within the bone marrow compartment, mainly residing along sinusoids (∼70% of MKPs) (**Fig. 1a** **and Extended Data Figs. 1b, c**).

**Figure 1.**
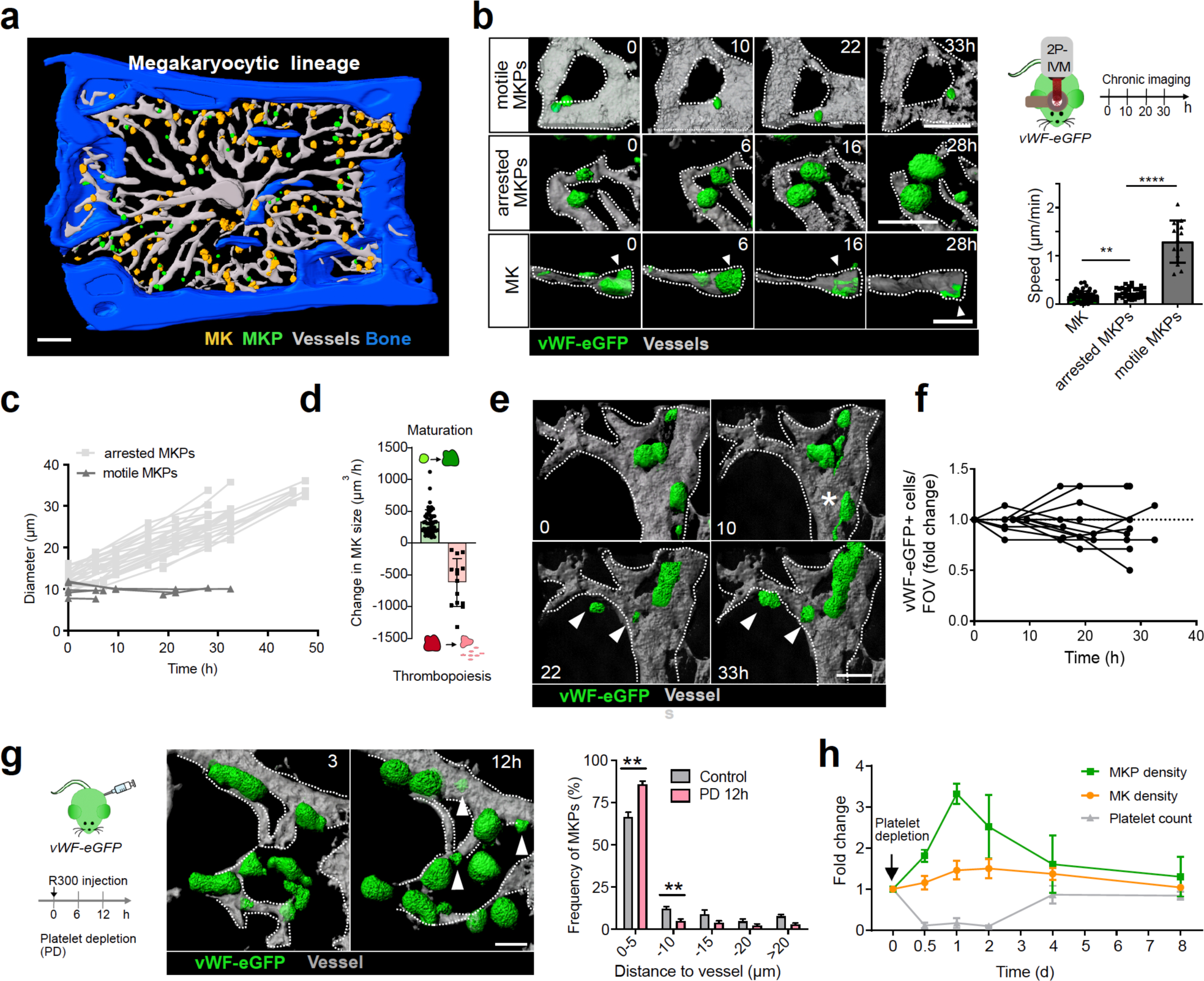
The megakaryocytic BM niche spatio-temporally coordinates megakaryopoiesis and thrombopoiesis. **a**, 3D rendered z-stack of murine BM (sternal bone). MKPs (green): CD41^+^/CD42^-^; MKs (yellow): CD41^+^/CD42^-^; blood vessels (grey): CD144^+^; bone (blue): second harmonic generation. **b**, Chronic 2P-IVM of calvarian bone; green: vWF^eGFP/+;^ grey: TRITC-dextran (blood vessels). Upper: MKP migrating at the perivascular niche; middle: MKPs lodged within the perivascular niche grow in size; lower: MK during thrombopoiesis, arrowheads indicate release of proplatelets. Single cell tracking revealed speeds of MKs (n=52), arrested MKPs (n=33) and motile MKPs (n=31); cells pooled from 7 mice, unpaired t-tests **: p=0.004; ****: p<0.0001. Mean±SEM. **c**, Diameters of single arrested and motile MKPs tracked over time (2P-IVM); arrested MKPs (n=27 cells from 5 mice); motile MKPs (n=6 cells from 3 mice). **d**, Change in cell volume per hour was measured for MKP maturation (growth; green) (n=14 cells from 4 mice) and MK thrombopoiesis (platelet release) (reduction; red) (n=11 cells from 4 mice); Mean±SEM **e,** Time series of 3D rendered z-stacks of calvarian BM. Arrow heads indicate growing MKPs. Asterisk highlights proplatelet forming MK. Notably, MK vanished at t=21.5h, while new MKPs appear within close proximity. **f**, Fold change of vWF^eGFP/+^ cells per field of view (FOV) tracked over time. n= 12 FOVs from 5 mice. **g**, Chronic 2P-IVM following PD (12h after injection of R300). Histogram shows increased frequency of MKPs within the perivascular niche after PD; n=4 mice per group **: p= 0.0011, and **: p=0.0062; Mean±SEM. **h**, Fold change of platelet counts (hemocytometer), MK and MKP density (number/mm^3^ marrow) (whole mount immunostainings) measured at indicated time points following platelet depletion; n≥3 mice. Scale bars= 50 µm.

To study the spatio-temporal patterns of megakaryopoiesis *in vivo* we performed 2P-IVM of *vWF-eGFP* reporter mice that specifically label the entire megakaryocytic lineage including MKPs and mature MKs^8^ (**Extended Data Figs. 1f-h, Methods)**. We visualized the same field of view for up to three days with a chronic imaging window implanted into the calvarial bone to track individual MKPs and MKs (**Extended Data Fig 2a**). We identified small, motile vWF^eGFP/+^ cells within the BM parenchyma that arrest along BM sinusoids **(Fig. 1b, Extended Data Fig. 2b)** and increase their volume by ∼10 fold (**Figs. 1c,d** **and Supplementary Video 2**), representing MKPs undergoing cytoplasmic maturation into large, sessile MKs (**Fig. 1b, Extended Fig. 1f)**. This provides first real-time evidence that perivascular positioning of immotile MKs is determined by their motile progenitors, as previously proposed by others^9^.

**Figure 2.**
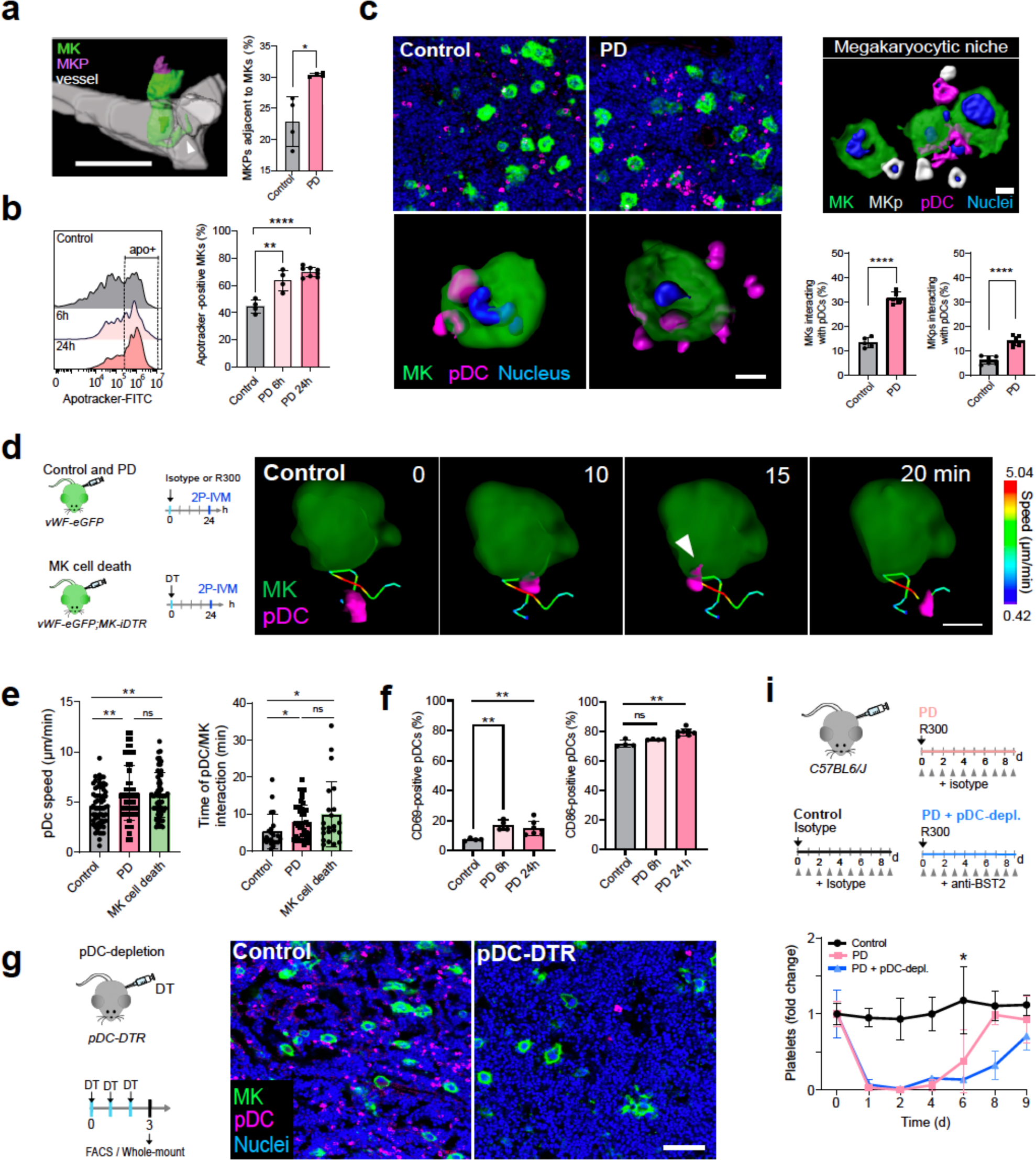
pDCs are BM niche cells that regulate megakaryopoiesis. **a,** Immune-mediated thrombocytopenia (PD) (also see Extended Data Fig. 3) triggers MKP proliferation in close contact to mature MKs. 3D rendered z-stack (BM whole-mount); MK (green): CD41^+^CD42^+^; MKP (purple): CD41^+^CD42^-^; vessels CD144^+^ (gray). Percentage of MKPs attached to MKs in control and following PD (12h); n=4 mice. Unpaired t-test/Welch’s correction; *: p=0.0316; Mean±SEM; Scale bar=100 µm. **b**, Frequency of apototic MKs increases following PD (pink: 6h; red: 24h) (FACS). MKs were stained and gated for CD41-PE^+^, CD42-APC^+^, CD11b^-^ CD8a^-^, Apotracker green^+^ and life/dead 405nm (dead cells exclusion). Left: Histograms of fluorescence intensity (Apotracker); right: frequency of apotracker-positive MKs; n≥4 mice; unpaired t-test; **: p=0.0058; ****: p<0.0001; Mean±SD. **c**, BM whole mounts show increased pDCs-MK/MKp interactions following PD; MKs (green, CD41^+^); MKPs (white, CD41^+^ and diameter > 15 µm); pDCs (purple, BST2^+^); nuclei (blue, DAPI). Scale bars = 10 µm. Bar plot: frequencies of pDC-MK- and pDC-MKP-interactions; n≥5 mice; unpaired Welch’s t-test; ****: p<0.0001; Mean±SD. **d**, 2P-IVM reveals migratory pattern of DCs and long-lasting pDC-MK interactions (calvarian BM). Schematic shows experimental design for **d** and **e**. MK: vWF-eGFP^+^ (green); pDCs: anti-siglecH-PE (i.v. 15min before imaging). pDCs protrude into MKs forming close contacts (arrowhead); color-code: pDC speed; scale bar=10 µm; also see Extended Data Fig. 5c. **e**, Single cell tracking of pDCs in control mice and after PD (R300) and induction of MK cell death (*PF4-Cre;RS26-iDTR; vWF-eGFP^+/-^*, also see Methods) reveals speed and times of MK-pDC interaction within the megakaryocytic niche. pDC speed: control: n=62 cells, PD: n= 60 cells, MK cell death: n= 62 cells, **: p=0.0030, **: p=0,0022, ns=0.644, pooled from 4 mice. Mean±SD. MK-pDC interaction: control: n=22 cells, PD n=42 cells, MK cells death: n=22 cells, pooled from 4 mice; unpaired t-test, *: p= 0.0418 and 0.0369; ns=0.3160. Mean±SD. **f**, FACS analysis of activation markers of BM pDCs following PD (pink: 6h; red: 24h). n= 4 mice; unpaired t test Welch’s correction; for CD69-postive: **: p=0.0081 and p=0.0055, for CD86-posotive **: p=0.0014, ns=0.0910; Mean±SD. **g-h**, Impaired megakaryopoiesis following pDC-depletion in *BDCA2-DTR (pDC-DTR)* mice (also see Methods). Cell numbers were quantified by histology (whole mount BM; MKs (green): CD41^+^; pDCs (magenta): BST2^+^) or FACS (see Extended Data Fig. 5d); platelets counts (hemocytometer). n ≥ 5 mice; unpaired t-test Welch’s correction; **: p=0.0012, ****: p <0.0001; Mean±SD. **i**, Schematic showing experimental design of platelet recovery experiment. Platelets were depleted by R300 on d0 and recovery of blood platelet counts was monitored at indicated time points for 9 days in the absence (antibody-mediated depletion: a-PDCA1 i.v. daily) and presence (isotype) of pDCs; n=3 mice; paired t-test; *: p=0.024; Mean±SD.

Mature MKs lodged within the perivascular niche release proplatelets to produce platelets (**Fig. 1b** **lower panel**)^3^. Once entering thrombopoiesis MKs rapidly reduce their volume and disappear completely within hours (**Fig. 1d** **and Extended Data Fig. 2c**). Consumption of platelet-producing MKs is irreversible as we did not observe recovery once MKs completed thrombopoiesis. Instead, new vWF^eGFP/+^ MKPs appear in proximity to vanished MKs giving rise to mature MKs (**Fig. 1e, Extended Data Fig. 2d and Supplementary Video 2**). Consequently, the total number of vWF^eGFP/+^ cells remains remarkably stable over several hours to days (**Fig. 1f**). Megakaryopoiesis and thrombopoiesis therefore are well synchronized processes that ensure immediate replenishment of platelet-producing MKs from their progenitors.

We tested whether megakaryopoiesis and thrombopoiesis also remain synchronized in situations of high platelet demand. We removed the entire circulating platelet pool by antibody-mediated platelet depletion (PD) (**see Methods, Extended Data Fig. 3a**). During PD, MKs lose sphericity indicating activation and engage in emergency platelet production through intrasinusoidal proplatelet extensions or MK fragmentation resulting in rapid reduction in MK size (**Extended Data Figs. 3b, c**). Chronic 4D-imaging revealed that the fast release of platelets from MKs is accompanied by an increased MKP proliferation that peaks at 12-24h after treatment and is most prominent at the perivascular niche (**Fig. 1g, and Extended Data Figs. 3d, e, Supplementary Video 2**). Accelerated proliferation of MKPs fully compensates for the high MK demand during emergency thrombopoiesis and replenishes the circulating platelet pool within 4 days while maintaining MK homeostasis (**Fig. 1h**). Megakaryopoiesis therefore is tightly coordinated to ensure MK homeostasis both in steady state and during pathological platelet consumption.

**Figure 3.**
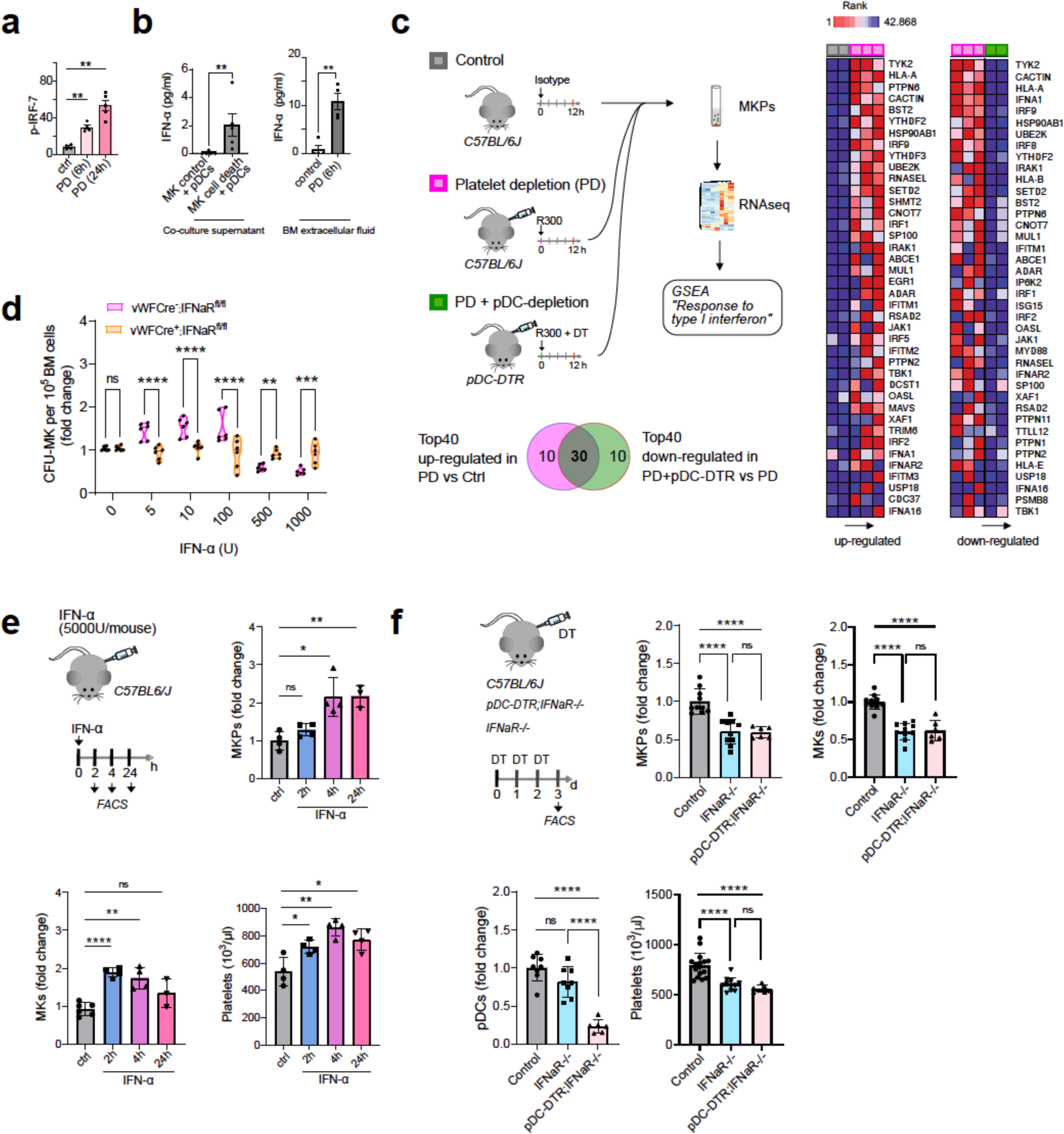
pDC-derived IFN-α fuels megakaryopoiesis. **a,** Elevated p-IRF-7 in pDCs following PD (FACS). n ≥ 4 mice; **: p=0.0011; Mean±SEM. **b,** Left: Co-culture of BM-derived pDCs and MKs. MK cell death was induced by DT injection of *PF4-Cre;RS26iDTR* mice; PBS-injected mice served as control. After 18h of co-culture IFN-α was measured in supernatants (ELISA); n=5 experiments; Mann Whitney test; **: p=0.0079; Mean±SD. Right: IFN-α was measured in BM extracellular fluid in control mice and following PD (6h) (ELISA); n= 5 mice; paired t-test; *: p=0.0317; Mean±SD. **c**, Schematic shows design of RNAseq experiment. MKPs were sorted from BM of mice treated with isotype antibody (grey; n=2), R300 (PD, purple; n=3) or R300 plus DT (PD plus pDC depletion, green; n=2). Gene set enrichment analysis was performed on RNA-Seq data. Heatmaps show differentially regulated genes. Heatmaps show the top 40 up- or down-regulated genes in the gene set “Response to type I interferon”, intersection analysis (Venn diagram) confirms a high overlap of genes (30 genes) to be inversely regulated between conditions. Also see Extended Fig. 7. **d**, MK-CFUs were analyzed in the presence or absence of IFN-α. Of note, while IFN-α concentrations ::100U increased MK-CFUs, concentration ≥ 500U showed adverse effects. Conditional depletion of IFNaR in megakaryocytic progenitors (*vWF-Cre;IFNaR^fl/fl^)* confirmed a direct and IFNaR-dependent role of IFN-α. n=6 experiments; two way ANOVA Multiple comparisons, Sidak’s multiple comparisons test,; ns= > 0.999, **: p=0.0061, ***: p=0.0003, ****: p=<0.0001 violin plot with Median (red line). **e**, Increased MKP, MK and platelet numbers in response to universal type I interferon alpha treatment (5000U, i.v.). MKPs, MKs (FACS of BM) and platelets (hemocytometer) were measured at indicated time points. n=3-4 mice; unpaired t-test; MKPs n= 3-4 mice; *: p=0.0137, p:**=0.0051, ns: p=0.181; MKs **: p=0,0055, ****: p<0.0001, ns=0.1758; platelets *: p=0.0320, **: p= 0.0034, *: p= 0.0132; Mean±SD. **f**, Decreased MKP, MK and platelet numbers in *IFNaR-/-* mice. Notably, depletion of pDCs in *pDC-DTR;IFNaR-/-* mice had no additive effect. pDCs, MKPs, MKs (FACS of BM) and platelets (hemocytometer); n=6-16 mice; paired unpaired t-test; ****: p<0.0001, ns=0.8576; Mean±SD. Also see Extended Data Fig. 7.

### pDCs are BM niche cells that regulate megakaryopoiesis

We next aimed to identify the mechanisms that control steady-state and emergency replenishment of MKs from proliferating MKPs. Thrombopoietin (TPO) is the most potent cytokine promoting megakaryopoiesis and plasma levels are tightly regulated by TPO sequestration on thrombopoietin receptors (cMPL) of circulating platelets^10^. However, TPO did not trigger local perivascular megakaryopoiesis (**Extended Data Figs. 4a-e**).

**Figure 4.**
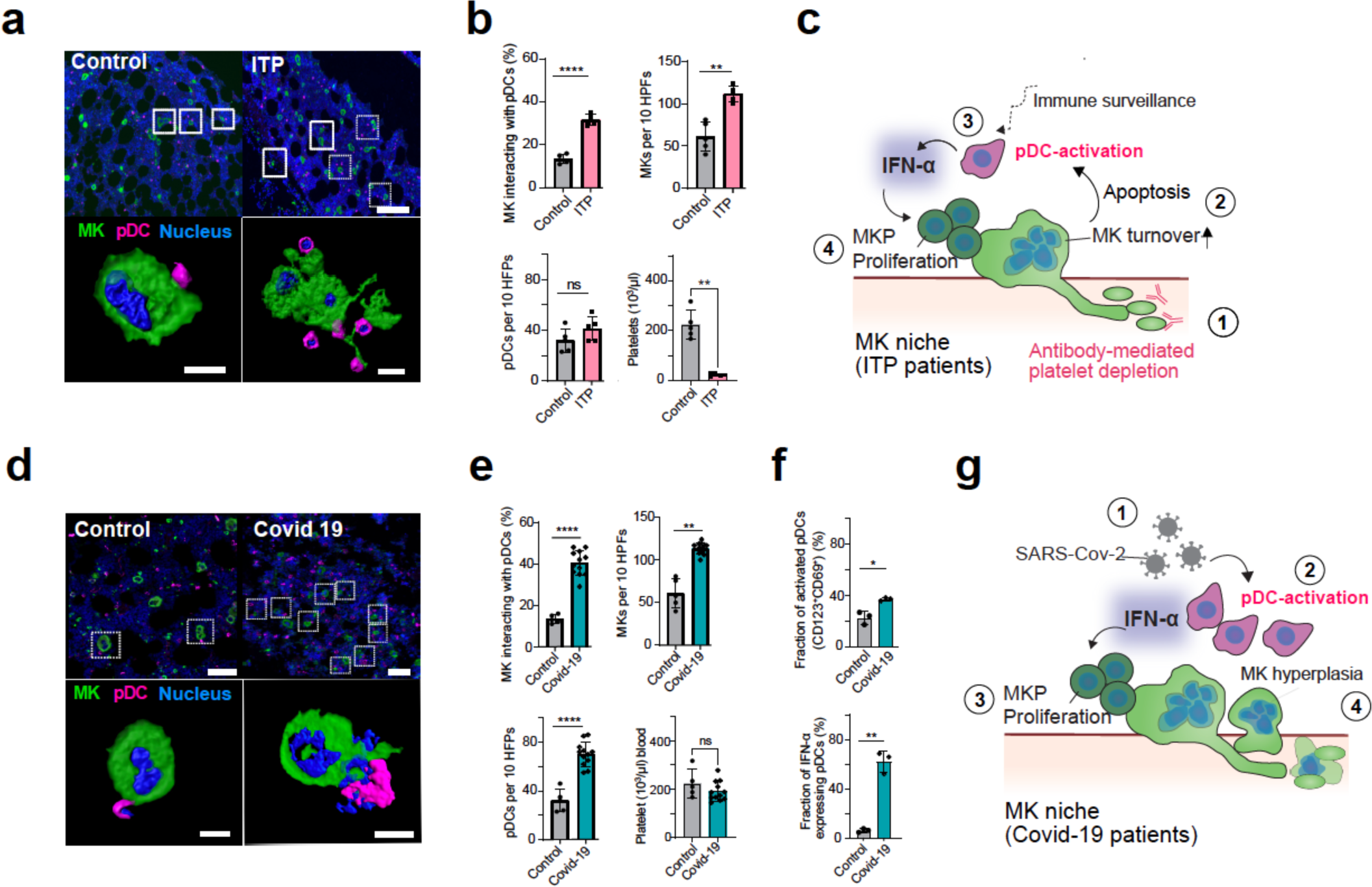
pDCs drive megakaryopoiesis in human BM. **a**, Immunohistology of human BM biopsies from healthy control and ITP patients. MK (CD41^+^, >15μm; green), pDCs (CD123^+^; magenta), nuclei (DAPI; blue). Lower micrographs: 3D reendered high magnification confocal z-stacks. **b**, Quantification of **a**, Cell numbers and interactions were measured in 10 HPFs (=…); platelet counts (hemocytometer); n=5 patients; unpaired t-test; MKs **: p=0.0011, MK/pDCs ****: p<0.0001; pDCs ns: p=0.1577, platelets **: p=0.0026; Mean±SD. **c,** Graphical summary: pDCs are sentinels of MK turnover (apoptosis) (2). Activated pDCs release INF-α within the MK niche (3) triggering MKP proliferation (4). Of note, while these processes occur in steady state they are boosted by an increased platelet demand as seen in ITP (1). **d,** Immunohistology of human BM biopsies from healthy control patients (same patients as shown in **a** and **b**) and from autopsies of Covid-19 patients (see **a**). **e,** Quantification of **d**, (see **b**,); n= 3 patients; unpaired t-test; MK/pDC ****: p<0.0001, MKs **: p= 0.0018, pDC ****:p<0.0001, platelet ns= 0.815; Mean±SD. **f,** Increased activation of pDCs within BM of Covid-19 patients. Quantification of activation marker CD69 (upper) and INF-α expression (lower) (Immunohistology). n= 3 patients; unpaired t-test; *: p=0.0106; **: p=0.0069; Mean±SD. **g**, Graphical summary of proposed pathomechanism: SARS-CoV2 hijacks BM pDCs which accumulate in the BM (1). Activation of pDCs and uncontrolled INF-α release (2) within the MK niche trigger MKP proliferation (3) and MK hyperplasia (4).

Whole-mount analysis of mouse BM shows that a considerable fraction of MKPs is located in close proximity to mature MKs (**Fig. 2a**)^11^. Based on this observation we hypothesized that vanishing MKs that release platelets and show signs of apoptosis^12^ may trigger their own replacement from local MKPs within the perivascular niche. When we triggered emergency thrombopoiesis by depleting platelets, the number of apoptotic MKs was increased (**Fig. 2b**). Different phagocyte subsets are equipped to sense and clear apoptotic bodies and DNA. In particular, macrophages play an important role in homeostasis of various tissues, including erythropoietic islands of the BM^13^. We found that approximately 12% of mature MKs colocalized with CD68+ macrophages during steady state (**Extended Data Fig. 5a**). However, these macrophage-MK interactions did not change during immune-mediated thrombocytopenia despite the increase in apoptotic MKs (**Fig. 2b, Extended Data Fig. 5a**) making their contribution to megakaryopoiesis less likely.

Plasmacytoid dendritic cells (pDCs) are another subset of immune cells that are specialized in detecting apoptotic cells and nuclei acids^14, 15^. While pDCs are rare in peripheral tissues, they are abundant in the BM, where they originate^16^. We show here that in mouse BM approximately 15% of mature MKs colocalize with BST2^+^ pDCs during steady state (**Fig. 2c**). These interactions increase threefold in response to antibody-mediated thrombocytopenia (**Fig. 2c**). Yet, the total number of pDCs in the bone marrow remained unaffected following platelet depletion, suggesting specific rather than stochastic recruitment to the megakaryocytic niche (**Fig. 2c**, **Extended Data Fig. 5b**). To analyze the dynamics of MK-pDC interactions *in vivo* we tracked fluorescently-labelled pDCs (anti-SiglecH-PE) in the calvarial bone marrow of vWF-eGFP mice (**Fig. 2d, Supplementary Video 3**). pDCs migrate randomly in the BM with mean speeds of 0.8µm/min (**Fig. 2e**). When pDCs encounter MKs, they decelerate and start crawling on the surface of MKs, often diving into the cell body (**Fig. 2d, Supplementary Video 3**), a process resembling emperipolesis^17^. To study if this scanning behavior changes in response to MK exhaustion, we induced cell death in megakaryocytes by treating *PF4-Cre;RS26-iDTR^fl/fl^;VWF-eGFP^+/-^* mice (referred to as MK-iDTR) with diphtheria-toxin (DT)^18, 19^ **(Extended Data Fig. 5c)**. MK cell death accelerated motility of pDCs (**Fig. 2e, Extended data Fig. 5c, Supplementary Video 3)**. At the same time, the duration of pDC-MK interactions increased (**Fig. 2e, Extended Data Fig. 5c**). Hence, pDCs monitor the BM by random walks and adapt their motility pattern when encountering dying MKs. Antibody-mediated thrombocytopenia triggered pDC activation (**Fig. 2f**) and phenocopied the motility patterns of pDCs observed in MK-iDTR-mice (**Fig. 2e, Extended Data Fig. 5c, Supplementary Video 3**).

To investigate whether MK-pDC interactions are essential to control megakaryopoiesis and MK homeostasis *in vivo* we depleted pDCs by treating BDCA2-DTR mice (pDC-DTR-mice) with DT^20^ (also see Methods). After 3 days of treatment approximately 80% of pDCs were depleted from the BM (**Fig. 2** **g, h**). Analysis of the megakaryocytic lineage by FACS and whole-mount immunostainings (**Fig. 2h**, **Extended Data Figs. 5d, 6)** revealed severely impaired megakaryopoiesis with a substantial decrease in MKPs (50%) and MKs (25%) in response to pDC depletion. Importantly, positioning of MKs and MKPs within the BM compartment was significantly altered compared to control mice indicating a crucial role of pDCs in maintaining the megakaryocytic niche (Extended Data **Fig. 5e**). Defective MK homeostasis in pDC ablated mice was also associated with thrombocytopenia with a 40% drop in circulating platelet counts in steady state (**Fig. 2** **h)**. In addition, in the setting of immune-mediated thrombocytopenia recovery of platelet counts was substantially delayed by more than 2 days and did not return to baseline in the absence of pDCs, indicating a reduced ability of pDC-depleted mice to cope with emergency thrombopoiesis (**Fig. 2i**). These findings identify pDCs as sentinel cells within the BM niche that sense exhausted (apoptotic) MKs and drive “on demand” megakaryopoiesis to secure MK and platelet homeostasis.

### pDC-derived IFN-α fuels megakaryopoiesis

pDCs encountering apoptotic cells release IFN-α in an IRF7-dependent manner thereby triggering local inflammation^16^. We found upregulation of phosphorylated IRF7 in pDCs following PD (**Fig. 3a**). Supernatants of pDCs co-cultured with apoptotic MKs contained robust amounts of IFN-α, while IFN-α was barely detectable in co-cultures with vital MKs (**Fig. 3b**). Similarly, IFN-α levels in the BM significantly increased in response to antibody-mediated thrombocytopenia (**Fig. 3b**) and both, MKs and MKPs expressed the IFN-α receptor (IFNaR) (**Extended Data Figs. 7a, b**). RNAseq analyses revealed that MKPs proliferating following PD (**Extended Data Figs. 7c-e**) significantly upregulate IFN-α-signaling and -response genes (**Fig. 3c, Extended Data Fig. 7f**). Depletion of pDCs in thrombocytopenic mice blunted upregulation of genes associated with proliferation and IFN-α, supporting pDCs as the major cellular source of IFN-α in this context (**Fig. 3c, Extended Data Fig. 7c-f**). IFN-α and TPO synergistically boosted megakaryopoiesis in an IFNaR-dependent manner *in vitro* (**Fig. 3d**). Moreover, systemic treatment of mice with IFNα triggered fast and immediate megakaryopoiesis *in vivo*, doubling MKP (**Fig. 3e**) as well as MK numbers within 2-4h (**Fig. 3e**) and increased circulating platelets by 1.5-fold (**Fig. 3e**). Similar to pDC-DTR mice, IFNaR-deficient animals showed reduced MKP numbers in steady state and were insufficient to maintain MK and platelet homeostasis (**Fig. 3f** and **Extended Data Fig. 7g**). IFNaR-/- BM-chimeras phenocopied global IFNaR-deletion (**Extended Data Fig. 7h)**. Ablation of pDCs in IFNaR-/- mice (*Clec4C-DTR; IFNaR-/-* mice) had no additive effect on either MKP, MK or platelet numbers (**Fig. 3f**). Hence, pDCs encountering exhausted/apoptotic MKs locally release IFN-α, which in turn drives expansion of MKPs via IFNaR-signaling to maintain MK homeostasis.

### pDCs drive megakaryopoiesis in human bone marrow

To define whether pDC-MK interactions are present in humans, we analyzed BM sections from healthy individuals (**Fig. 4a**). Consistent with our findings in mice, approximately 12% of MKs co-localized with pDCs under steady state (**Fig. 4a, b**). We then examined the BM of a cohort of patients with severe immunothrombocytopenia (ITP) (**see Supplementary Table for patient characteristics**). Similar to PD in mice, the number of circulating platelets was severely reduced in ITP patients (**Fig. 4b**). Accordingly, pDC-MK interactions were 3-fold higher in ITP compared to control patients while the number of MKs doubled (**Fig. 4b**). This suggests that pDCs are sentinels of MK turnover also in human BM (**Fig. 4c**).

pDCs are specialized to fight viral infections and play a key part in antiviral immunity^21, 22^. They have been involved in the control of coronavirus infection including the ongoing pandemic of coronavirus disease 2019 (COVID-19) caused by severe acute respiratory syndrome coronavirus 2 (SARS-CoV-2)^23, 24^. pDCs are activated by SARS-CoV-2 and contribute to type I IFN-dependent immunity during COVID-19^25^. pDC numbers in peripheral blood of COVID-19 patients correlate strongly with disease severity^26^. At the same time, increased numbers of naked megakaryocyte nuclei have been described in the BM and lungs from patients with severe COVID-19, that were linked to thrombotic complications^27^. To address whether SARS-CoV-2 infection is associated with dysregulation of pDC-driven megakaryopoiesis we analyzed BM of humanized mice susceptible to SARS-CoV-2 (*FVB-K18hACE2*). 6 days post infection mice showed elevated pDC numbers within the BM. pDCs engaged in close interaction with MKs, and both MKP and MK numbers increased compared to uninfected controls suggesting pDC-driven megakaryopoiesis (**Extended Data Fig. 8**). We next analyzed human BM from a cohort of patients that died from COVID-19 (**see Supplementary Table for patient characteristics**). We found a more than 2-fold increase in MK and pDC numbers, and a 3-fold increase in MK-pDC interactions in COVID-19 patients compared to controls (**Figs. 4d, e**). pDCs respond to SARS-CoV-2 by producing type I interferons^28^ and severe COVID-19 is associated with an expansion of interferon-activated MKs in the circulating blood^29^. Accordingly, the fraction of activated (CD69+) and IFN-α-expressing pDCs in COVID-19 BM was significantly increased compared to healthy controls **(Fig. 4f, g).** This suggests that pDC-derived type I interferon drives megakaryocytic hyperplasia in COVID-19. Excessive activation of pDC-induced megakaryopoiesis could therefore contribute to dysbalanced MK homeostasis reported in severe COVID-19 and potentially also in the context of other viral infections.

## Discussion

Type 1 interferons are mainly recognized for their protective role in viral infections^30^. However, numerous physiological processes beyond antiviral defense, have been identified to rely on IFNs including immunomodulation, immunometabolism^31^, cell cycle regulation, cell survival and cell differentiation^31, 32^. Here, we show that IFN-α released by pDCs is the key signal to accomplish precise control of BM megakaryopoiesis and to ensure blood platelet homeostasis.

Within the BM compartment, constitutive IFN-α levels are required for maintenance of the hematopoietic stem cell (HSC) niche, but long-term systemic elevation of IFN-α levels may cause exhaustion of HSCs^33, 34^. In addition, long term treatments with high-dose IFN, as well as infections associated with persistently high IFN-α levels are associated with impaired platelet production and platelet function^35–39^. This indicates that IFN-α levels must be precisely controlled in a spatio-temporal fashion. We show here that pDCs are motile carriers of IFN-α that constantly patrol the BM and locally dispatch IFN-α when encountering apoptotic MKs that are about to quit platelet production. This identifies pDCs as critical element of the megakaryocytic niche, that control homeostasis of megakaryocytes and blood platelets through type 1 interferon signals that remain confined to the BM.

The control of megakaryocyte numbers by pDCs may serve functions beyond platelet homeostasis: The BM microenvironment is functionally compartmentalized by a heterogeneous population of niche cells that provide physical and soluble signals to spatio-temporally organize hematopoiesis^40^. Besides giving birth to platelets, MKs constitute niche cells of hematopoietic origin that regulate HSCs during homeostasis and stress^18, 19^. Consequently, maintenance of the megakaryocytic HSC niche requires seamless replenishment of consumed, platelet-producing MKs. We establish pDCs as an additional player in the niche orchestrating thrombopoiesis and megakaryopoiesis to maintain MK homeostasis and may thus also contribute to the maintenance of the megakaryocytic HSC niche^18, 19^.

Acute systemic inflammation is known to instruct emergency megakaryopoiesis to maintain and regenerate the platelet pool^37^. Our findings also support a role of pDCs in emergency megakaryopoiesis induced by systemic inflammation: In the perivascular MK niche, BM pDCs are in a strategic position to integrate systemic inflammatory signatures and translate them into enhanced platelet production. Platelets prevent blood loss, but also counteract infectious agents through interaction with neutrophils^41, 42^. Therefore, pDC-driven inflammatory megakaryopoiesis likely is beneficial in any type of acute tissue injury associated with loss of vascular integrity and platelet consumption. However, it may also be detrimental, when pDC-driven megakaryopoiesis is mismanaged, e.g. during severe viral diseases. A case in point is infection with corona virus SARS-CoV-2, which dysregulates the fine-tuned IFN-α production of pDCs^43^. While analysis of peripheral blood of SARS-CoV-2 infected patients revealed reduced pDC counts with muted IFN-α production^43^, we identify an accumulation of activated, IFN-α releasing pDCs in the BM of patients with severe disease progression. pDCs engage in close contact to MKs which is accompanied by marked megakaryocytic hyperplasia. Since expansion of IFN-α-activated MKs is a hallmark of severe Covid-19^29^ and associated with thrombocytopathy, thrombosis and organ failure^6^, our data identifies SARS-CoV-2-hijacked pDCs as promising therapeutic target to restore MK homeostasis to prevent detrimental thrombotic insults. In conclusion, we identified a novel role of pDCs in orchestrating blood platelet homeostasis. Targeting pDC-driven megakaryopoiesis opens new options to boost or suppress platelet production in different clinical scenarios.

## Methods

### Materials

| REAGENT or RESOURCE | SOURCE | IDENTIFIER |
| --- | --- | --- |
| **Antibodies and dyes** |  |  |
| Ultra-LEAFTM purified anti-PDCA-1, clone 927 | BioLegend | Cat# 114110 |
| Ultra-LEAF™ Purified Rat IgG2b, κ Isotype Ctrl (clone RTK4530) | BioLegend | Cat# 400671 |
| Human IFN Alpha Hybrid Protein (Universal Type I IFN) | PBL, assay science | Cat# 11200-1 |
| Anti-human CD123 | Abcam | Cat# ab205365 |
| Anti-CD41 Recombinant (EPR4330) | Abcam | Cat# ab134131 |
| Anti-human CD69 Monoclonal (8B6) | ThermoFisher | Cat#MA5 15612 |
| Anti-IFN alpha Polyclonal Antibody | ThermoFisher | Cat#PA5-115430 |
| Donkey anti-Rabbit IgG (H+L) Highly Cross-Adsorbed Secondary Antibody, Alexa Fluor 488 | ThermoFisher | Cat# A-21206 |
| Donkey anti-Mouse IgG (H+L) Highly Cross-Adsorbed Secondary Antibody, Alexa Fluor 555 | ThermoFisher | Cat# A-31570 |
| Donkey anti-Goat IgG (H+L) Cross-Adsorbed Secondary Antibody, Alexa Fluor 647 | ThermoFisher | Cat# A-21447 |
| Purified anti-mouse/rat CD42d Antibody (clone 1C2) | BioLegend | Cat# 148502 |
| Goat Anti-Armenian hamster IgG H&L Secondary Antibody, Alexa Fluor 647 | Abcam | Cat# ab173004 |
| FITC anti-mouse CD41a (clone MWReg30) | eBioscience | Cat# 11-0411-85 |
| APC anti-mouse/rat CD42d (clone 1C2) | BioLegend | Cat# 148506 |
| Biotin anti-mouse VE-cadherin (clone eBioBV13) | eBioscience | Cat# 13-1441-82 |
| Streptavidin PE | eBioscience | Cat# 12-4317-87 |
| SiglecH anti-mouse Monoclonal Antibody (eBio440c), FITC | eBioscience | Cat# 11-0333-82 |
| SiglecH anti-mouse Monoclonal Antibody (eBio440c), PE | eBioscience | Cat# 12-0333-82 |
| SiglecH anti-mouse Antibody, PerCP/Cyanine5.5 | BioLegend | Cat# 129614 |
| CD317 (BST2, PDCA-1) anti-mouse Antibody, APC | BioLegend | Cat# 127016 |
| CD317 (BST2, PDCA-1) anti-mouse monoclonal Antibody (eBio927), Functional Grade | eBioscience | Cat# 16-3172-81 |
| Goat Anti-Rat IgG H&L, Secondary Antibody, Alexa Fluor 647 | Abcam | Cat# ab150159 |
| Platelet depletion antibody isotype control | emfret | Cat# C301 |
| Platelet depletion antibody | emfret | Cat# R300 |
| PE anti-human CD41 antibody (clone PM6/248) | Biorad | Cat# MCA467PE |
| TER-119 Monoclonal Antibody (TER-119), Biotin | eBioscience | Cat # 13-5921-82 |
| CD3 Monoclonal Antibody (OKT3), Biotin | eBioscience | Cat # 13-0037-82 |
| CD45R (B220) Monoclonal Antibody (RA3-6B2), Biotin | eBioscience | Cat # 13-0452-82 |
| CD11b Monoclonal Antibody (M1/70), Biotin | eBioscience | Cat # 13-0112-82 |
| Ly-6G Monoclonal Antibody (1A8-Ly6g), Biotin | eBioscience | Cat # 13-9668-82 |
| Pacific Blue™ anti-mouse TER-119 Antibody | BioLegend | Cat # 116232 |
| Pacific Blue™ anti-mouse CD3 Antibody | BioLegend | Cat # 100214 |
| Pacific Blue™ Rat Anti-Mouse CD45R Antibody | BD bioscience | Cat # 558108 |
| Pacific Blue™ anti-mouse/human CD11b Antibody | BioLegend | Cat # 101224 |
| Pacific Blue™ anti-mouse Ly-6G Antibody | BioLegend | Cat # 127612 |
| Goat anti-Mouse IgG (H+L) Highly Cross-Adsorbed Secondary Antibody, Alexa Fluor 555 | ThermoFisher | Cat # A-21424 |
| IRF7 Monoclonal Antibody (MNGPKL), PE | eBioscience | Cat # 12-5829-82 |
| APC anti-mouse CD117 (c-Kit) Antibody | BioLegend | Cat # 105812 |
| PE/Cyanine7 anti-mouse Ly-6A/E (Sca-1) Antibody | BioLegend | Cat # 108114 |
| PE/Cyanine7 anti-mouse CD105 Antibody | BioLegend | Cat # 120410 |
| Brilliant Violet 510™ anti-mouse CD150 (SLAM) Antibody | BioLegend | Cat # 115929 |
| PerCP/Cyanine5.5 anti-mouse CD9 Antibody | BioLegend | Cat # 124818 |
| PE/Cyanine7 anti-mouse/human CD11b Antibody | BioLegend | Cat # 101216 |
| APC Rat Anti-Mouse CD45R/B220 | eBioscience | Cat # 553092 |
| FITC anti-mouse CD69 Antibody | BioLegend | Cat # 104506 |
| PE anti-mouse CD86 Antibody | BioLegend | Cat # 105008 |
| APC/Cyanine7 anti-mouse/human CD11b Antibody | BioLegend | Cat # 101226 |
| Dextran, Tetramethylrhodamine, 10,000 MW, Lysine Fixable (fluoro-Ruby) | ThermoFisher | Cat # D1817 |
| Dextran, Cascade Blue™, 10,000 MW, Anionic, Lysine Fixable | ThermoFisher | Cat # D1976 |
| Hoechst 33342, Trihydrochloride, Trihydrate | ThermoFisher | Cat# H3570 |
| DAPI (4',6-Diamidino-2-Phenylindole, Dihydrochloride) | ThermoFisher | Cat# D1306 |
| LIVE/DEAD™ Fixable Aqua Dead Cell Stain Kit, for 405 nm excitation | ThermoFisher | Cat# L34957 |
| SYTOX™ Orange Dead Cell Stain, for flow cytometry | ThermoFisher | Cat# S34861 |
| Apotracker™ Green | BioLegend | Cat# 427402 |
| Purified Rat Anti-Mouse CD16/CD32 (Mouse BD Fc Block™) | BD Bioscience | Cat# 553142 |
| **Reagents, drugs and recombinant proteins** |  |  |
| Diphtheria Toxin | Sigma-Aldrich | Cat# 322326 |
| Mouse recombinant thrombopoietin | Immunotools | Cat# 12343615 |
| Tissue Tek | Sakura Finetek |  |
| Click-iT™ EdU Alexa Fluor™ 647 Flow Cytometry Assay Kit | ThermoFisher | Cat# C10419 |
| Recombinant Mouse FLT3L (carrier-free) | BioLegend | Cat# 550706 |
| MegaCult™-C Complete Kit Without Cytokines | StemCell | Cat # 04900 |
| Collagen Solution | StemCell | Cat # 04902 |
| Mouse Thrombopoietin Quantikine ELISA Kit | R&D systems | Cat # MTP00 |
| Mouse IFN Alpha All Subtype ELISA Kit, High Sensitivity (Serum, Plasma, TCM) | PBL, assay science | Cat # 42115-1 |
| RNeasy Micro Kit | QIAGEN | Cat # 74004 |
| High-Capacity cDNA Reverse Transcription Kit | Applied Biosystems | Cat # 4368814 |
| SsoAdvanced Universal SYBR Green Supermix | Biorad | Cat # 172-5271 |
| Lysing Buffer | BD Bioscience | Cat # 555899 |
| EasySep™ Mouse CD19 Positive Selection Kit II | STEMCELL | Cat # 18954 |
| EasySep™ Mouse CD11b Positive Selection Kit II | STEMCELL | Cat # 18970 |
| Normal Goat Serum (10%) | ThermoFisher | Cat # 50197Z |
| Dako Fluorescence Mounting Medium | DAKO | Cat # S3023 |
| Cyano Venner dental fast glue | Hager Werken | Cat # 152261 |
| 123count eBeads™ Counting Beads | ThermoFisher | Cat # 01-1234-42 |
| Gibco™ RPMI 1640 Medium, GlutaMAX™ Supplement | Fisher scientific | Cat # 11554516 |
| Penicillin-Streptomycin-Glutamine (100X) | ThermoFisher | Cat# 10378016 |
| Sodium Pyruvate (100 mM) | ThermoFisher | Cat# 11360070 |
| MEM Non-Essential Amino Acids Solution (100X) | ThermoFisher | Cat# 11140035 |
| 2-Mercaptoethanol (50 mM) | ThermoFisher | Cat# 31350010 |

#### Mouse strains

*C57BL/6J, PF4-Cre (C57BL/6-Tg(Pf4-icre)Q3Rsko/J)*^44^*, Rosa26-iDTR^flox^ (C57BL/6^Gt^(ROSA)26^Sortm^*^1^*(HBEGF)^Awai/J^)*^45^, *IFNαR-/-* (*B6.129S2-Ifnar1^tm1Agt/Mmjax^*)^46^, *IFNαR1^flox^ (B6(Cg)-Ifnar1^tm^*^1^*^.1Ees^/J)*^47^ and *BDCA2-DTR* (*C57BL/6-Tg(CLEC4C-HBEGF)956^Cln/J^)*(referred to as *pDC-DTR*)^48^ mice were purchased from The Jackson Laboratory. *vWF-Cre* mice were generated by W. Aird and described previously^49^. *vWF-eGFP* mice were generated by C. Nerlov and described previously^8^. *PF4-Cre* were crossed with *Rosa26-iDTR* mice to induce megakaryocyte cell death *in vivo (PF4-Cre; RS26-iDTR)*^18^. *PF4-Cre;RS26-iDTR* were crossed with *vWF-eGFP* to visualize the megakaryocytic lineage following induction of MK cell death (referred to as *MK-iDTR*). *vWF-Cre* mice were crossed with *IFNαR1^flox^*to conditionally delete IFNaR in the megakaryocytic lineage. *pDC-DTR* and *IFNαR-/-* were cross bred to achieve pDC depletion in IFNaR-/- animals (*pDC-DTR; IFNaR-/-*).

Both male and female mice were used in this study. If not otherwise stated, animals of control and experimental group were sex-matched and age-matched (6–12 weeks). Animals were bred and maintained in the animal facilities of the Walter-Brendel Zentrum, the Zentrum für Neuropathologie und Prionforschung (ZNP) or the Biomedical center of the LMU Munich, all in Munich, Germany. All mice live in standardized conditions where temperature, humidity, and hours of light and darkness are maintained at a constant level all year round. All animal experiments were performed in compliance with all relevant ethical regulations for studies involving mice and were approved by the local legislation on protection of animals (Regierung von Oberbayern, Munich).

#### Mouse anesthesia

If not stated otherwise, anesthesia was performed by isoflurane induction, followed by intraperitoneal injection of medetomidine (0.5 mg/kg body weight), midazolam (5 mg/kg body weight), and fentanyl (0.05 mg/kg body weight). Toe pinching reflexes and breathing pattern were used to determine the adequate depth of anesthesia. Core body temperature was maintained by heating pads, and narcosis was maintained by repetitive injections of 50% of the induction dose, if necessary.

#### Human samples

LMU Munich: Bone marrow specimens of five patients with clinically proven ITP, of five patients with non-Hodgkin lymphoma without bone marrow involvement and 12 patients who died of COVID-19 were analyzed. The specimens of ITP and lymphoma patients were archived material, the COVID-19 specimens were taken during autopsy.Clincal details are given in supplemental table 1. The study was approved by and conducted according to requirements of the ethics committees at the Ludwig Maximilians University of Munich (20-1039). There was no commercial support for this study. University Clinic Aachen: We included 6 consecutive clinical autopsies of COVID-19 positive patients between March 9th, 2020 and May 5th, 2020 performed at the Institute of Pathology of the University Clinic Aachen. Each patient had a positive clinical SARS-CoV-2 PCR test from upper or lower respiratory tract prior to autopsy, confirmed by post-mortem RT-PCR. Consent to autopsy was obtained by the legal representatives of the deceased patients. The study was approved by the local ethics committee (EK 304/20, EK 119/20, and EK 092/20). Bone marrow samples were obtained with an electric autopsy saw (Medezine 5000, Medezine, Sheffield, UK) from the vertebral bodies. The autopsies were performed in two steps according to a modified standard protocol to further increase employee safety and sample acquisition (developed in the frame of the German Registry of COVID-19 autopsies, www.DeRegCOVID.ukaachen.de). Samples were decalcified in formic acid or EDTA prior to dehydration and embedding in paraffin. Formalin-fixed, paraffin embedded bone marrow blocks were cut on a microtome at 1–3 µm thickness and decalcified again in EDTA en bloque if necessary.

#### Drug treatments

Diphtheria Toxin (DT) was purchased from Sigma-Aldrich (322326, Saint Louis, MO, USA) and was injected intraperitoneally into *pDC-DTR*- and *pDC-DTR-IFNαR-/-* mice with a dose of 8 ng/g per day for consecutive 3 days. A single dose was injected into MK-iDTR mice 24h before the experiment. Platelet depleting antibodies (R300, anti-GPIbα) and isotype control (C301) were purchased from Emfret (Eibelstadt, Germany) and used according to the manufacturer’s protocol. pDC depleting antibody (ultra-LEAF^TM^ purified anti-PDCA-1, clone 927, BioLegend) was injected i.p. for 9 consecutive days with a concentration of 150 µg per mouse at day 1 and 100 µg per mouse the following days. The isotype control (ultra-LEAF^TM^ purified rat IgG2bk Isotype Ctrl, clone RTK4530, BioLegend) was injected accordingly. Type I interferon alpha was applied by injecting universal interferon alpha (PBL, assay science) with 5000U / mouse i.p in 200 µl PBS.

#### Bone marrow transplantation

Lin^-^ Sca-1^+^ c-kit^+^ (LSK) cells were isolated and sorted from bone marrow of *IFNαR-/-* and control mice (Lin-Pacific Blue (Ter-119, CD3, CD8a, CD45R, CD11b, Ly-6G), Sca-1-PE-Cy7, c-kit-APC, all purchased from Biolegend (San Diego, USA)). 8 × 10^3^ LSK cells were intravenously injected into lethally irradiated *C57BL/6J* female mice (two doses of 6.5Gy with a time interval of 8 hours). The bone marrow of chimeras was analyzed 8 weeks after the transplantation.

#### Mouse model of SARS-CoV2 infection. Mouse model of SARS-CoV2 infection

B6.Cg-Tg(K18-ACE2)^2Prlmn/^J mice (on a C57BL/6 background) were purchased from The Jackson Laboratory and bred against FVB mice to obtain C57BL/6 x FVB F1 hybrids. Mice were housed under specific pathogen-free conditions and heterozygous mice were used at 6-10 weeks of age. All experimental animal procedures were approved by the Institutional Animal Committee of the San Raffaele Scientific Institute and all infectious work was performed in designed BSL-3 workspaces. Mice were infected intranasally with 10^5^ TCID_50_ of SARS-CoV2 (hCoV-19/Italy/LOM-UniSR-1/2020 (EPI_ISL_413489) isolate) in 25μl. Six days after infection mice were perfusion fixed with 4% PFA and femurs were embedded in Tissue Tek (also see below). The frozen femurs were cut until the marrow was exposed. The femurs were rinsed with PBS and post fixed with 4% PFA for 15 min at RT. Femurs were washed with PBS and incubated with 10% goat serum for 1-2h at RT. Bone marrow was stained with anti-mouse CD41 (for MK/MKP), anti-mouse BST2 (for pDC), and Dapi for nucleus staining.

#### Immunohistology of human bone marrow samples

Bone marrow biopsies of five patients with confirmed immune thrombocytopenia and platelet counts < 30 x 10^9^/l were compared with age-matched controls (normal bone marrow biopsies performed for lymphoma staging). Tissue was fixed for 12 hours in 4% formalin and embedded in paraffin. For immunohistochemistry 1.5 µm sections were used. Multiplex-immunofluorescence or confocal laser scanning microscopy imaging were performed after antigen retrieval with epitope retrieval buffer (PerkinElmer Inc.). Slides were incubated sequentially for 1h with the following antibodies: pDCs: anti-human CD123 (#ab205365 Abcam 1:100); MKs: anti-human CD41 (#ab134131 Abcam 1:100) and detection was carried out by TSA-Opal620 (PerkinElmer) and TSA-Opal650 (PerkinElmer) or LSM 880 confocal microscope (Carl Zeiss). Multispectral imaging was completed using the PerkinElmer Vectra Polaris platform. Images were analyzed with HALO (Indica labs) software. In addition, samples were imaged with a LSM 880 confocal microscope using the Airyscan module (Carl Zeiss), Plan-Apo 20x/0.8 or 63x/1.46 objectives and analyzed with Zen Blue software (Carl Zeiss).

Bone marrow autopsies from Covid 19 patients (embedded in paraffin) were deparaffinized with xilol 2 x for 5 min, EtOH 100% 2 x for 2 min, EtOH 96% 1 x 3 min, EtOH 70% 1 x 2 min, and submitted to antigen retrieval with Tris-EDTA pH9 for 20 min, washed in 0.5 %BSA-PBS-Tween20 (0.1%) 1 x 5 min. Samples were blocked in 10% donkey serum with 0.5% saponin for 1 h RT. To monitor pDC activation and IFN alpha production, the following primary antibodies were used: CD69 mouse anti-human (#MA5-15612 Thermo Fisher), IFN-alpha rabbit polyclonal (#PA5-115430 ThermoFisher). CD123-goat polyclonal (#ab205365 Abcam) was used to label pDCs. Primary antibodies were incubated at 4 °C overnight and samples were subsequently washed with 0.5%BSA-PBS-Tween20 (0.1%) 3 x 5 min before adding secondary antibodies. Secondary antibodies (1:200): donkey anti-rabbit-AF488, donkey anti-mouse-AF 555, and donkey anti-gost-AF647. DAPI was used for nucleus staining (15 Min at RT). The samples were washed with PBS 3 x 5 min, before mounting with DAKO mounting medium.

#### Immunohistology of mouse bone marrow whole mounts

Mice were sacrificed and perfused fixed via the left ventricle (3 ml phosphate buffer saline (PBS) followed by 3 ml 4% paraformaldehyde (PFA)). Bones (sternum, femur and tibiae) were harvested and post-fixed in 4% PFA for 30 min at room temperature (RT), and incubated in 15% Sucrose for 2 hours at 4 °C and in 30% Sucrose at 4°C overnight. Afterwards, the bones were embedded in Tissue-Tek® O.C.T. Compound (Staufen, Germany) and frozen and stored at -80 °C. Frozen bones were cut on Histo Serve NX70 cryostat (Celle, Germany) until the exposure of the bone marrow. The sternum was cut as sagittal section. The femurs and tibiae were cut as coronal section or cross section, according to purpose. Bones were carefully removed from O.C.T. and gently washed in 1×PBS. For whole-mount staining, the cut bones were first incubated in 10% Normal Goat Serum (NGS, Thermo Fisher Scientific, Massachusetts, USA) and 0.5% Triton X-100 (Sigma-Alrich, Saint Louis, MO, USA) for at least 45min at RT (blocking/permeabilization). Bones were then incubated with primary antibodies at RT overnight and washed with PBS before adding secondary antibodies for 2 hours at RT. Labelling of MKs/MKPs: primary: CD41-FITC^+^ and CD42-purified hamster anti-mouse (BioLegend); secondary: goat anti-hamster Alexa Fluor 647 (Abcam) (1:100). Labelling of vessels: primary anti-VE-Cadherin (CD144) biotin purified (1:100); secondary: Streptavidin-PE (eBioscience) (1:200). Labelling of pDCs: primary: anti-SiglecH-PE/FITC or BST2 (CD317/PDCA-1-ThermoFisher-eBioscience) rat anti-mouse purified; secondary: goat anti-rat Alexa fluor 647. To label the nucleus Hoechst 33342 or DAPI, 1:1000 (ThermoFisher) was used. Lineage-biotin antibodies (Ter-119, CD3e, CD45R, CD11b, Ly-6G,) and streptavidin-PE were used with dilution of 1:200, all antibodies were purchased from eBioscience (San Diego, USA). IFN alpha staining: primary: IFN alpha polyclonal antibody (PA5115430) (1:100); secondary: goat-anti-mouse rabbit 555 (1:200) (ThermoFisher Scientific). After staining, bone samples were imaged using a multiphoton LaVision Biotech (Bielefeld, Germany) TrimScope II system connected to an upright Olympus microscope, equipped with a Ti;Sa Chameleon Ultra II laser (Coherent) tunable in the range of 680 to 1080 nm and a 16 × water immersion objective (numerical aperture 0.8, Nikon). Single images were acquired in 50-80µm depth, with z-interval of 2µm. The signal was detected by photomultipliers (PMTs) (G6780-20, Hamamatsu Photonics, Hamamatsu, Japan). ImSpector Pro (LaVision) was used as acquisition software.

In addition, bones were imaged with a LSM 880 confocal microscopy using the Airyscan module, objective Plan-Apo 20x Objective NA, 0.8 or with 63x/1.46 oil Plan-Apo. Images were taken with step size of 2 µm, range in z-stack of 40 µm, and analyzed with Zen Blue software. 3D projections and rendering were done with Imaris 9.2 (Bitplane) software.

#### Multiphoton intravital imaging of the calvarian bone marrow

Anesthetized mice were placed on a metal stage with a warming pad to maintain body temperature. The hair over the skull was carefully removed with an electric hair clipper. The skin on the skull was then cut in the midline to expose the frontal bone. For short term imaging (<4h), a custom-build metal ring was glued directly on the center of the skull, and the mouse’s head was immobilized by fixing the ring on a stereotactic metal stage. After imaging the mice were euthanized by cervical dislocation. For long term (chronic) imaging, a chronic window was implanted on the skull. Briefly, a round cover glass (diameter: 6 mm) was centered on top of the frontal bone with sterile saline in between glass and the bone surface. The surrounding area of the glass was then filled with dental glue (Cyano veneer, Hager Werken, Germany) and a custom plastic ring with inner diameter 8 mm was carefully centered on the frontal bone, with the glass exactly in the middle of the ring. The ring was further immobilized by applying the glue in the gap between the outer edge of the glass and the inner edge of the ring, as well as the gap between the outer edge of the ring and the tissue. Surgery was performed under sterile conditions. The mouse calvarium was imaged using a multiphoton LaVision Biotech (Bielefeld, Germany) TrimScope II system connected to an upright Olympus microscope, equipped with a Ti;Sa Chameleon Ultra II laser (Coherent) tunable in the range of 680 to 1080 nm and additionally an optical parametric oscillator (OPO) compact to support the range of 1000 to 1600 nm and a 16 × water immersion objective (numerical aperture 0.8, Nikon). Timelapse movies of 3D stacks were recorded within 30 µm to 40 µm depth, with z-interval of 2 or 3 µm and a frame rate of 1 min. Chronic imaging was performed at frame rates < 6h. Blood vessels and bone structure were taken as landmarks to retrieve the same imaging area of the bone marrow. 3D z-stacks were acquired with z-interval of 2 µm. 870 nm or 900 nm was used as an excitation wavelength. The signal was detected by Photomultipliers (G6780-20, Hamamatsu Photonics, Hamamatsu, Japan). ImSpector Pro (LaVision) was used as acquisition software. Imaging was performed at 37℃ using a customized incubator. Blood vessel were visualized by i.v. injection of Dextran Tetramethylrhodamine (TRITC-Dextran, 100µg in 100µl solution, ThermoFisher Scientific) or Dextran Cascade Blue 10.000 MW (D1976, ThermoFisher Scientific) before imaging. *vWF-eGFP* mice was used to visualize the megakaryocytic lineage; pDCs were labeled with SiglecH-PE antibody (eBioscience) injected intravenously 20 min before imaging (20 µl diluted with 100 µl NaCl).

#### Image processing

Movies and images were analyzed with Imaris software version 9.2 (Bitplane) or ZEN blue software (Carl Zeiss). Mosaic images were stitched in Imaris. Numbers of MKs, MKPs and pDCs were quantified in the whole mosaic images and normalized by the total volume of the BM in the image. The cell distance to vessels and/or endosteal surface was measured manually in Imaris Slice mode or by using ZEN blue software. The mean diameter of a MKP/MK was calculated by the average of the longest and shortest axis of the cell. Cell volumes of 3D rendered bone marrow stacks were measured automatically in Imaris. Cell migration was analyzed in 3D time-lapse movies by tracking the cell at every time point (Imaris). The cell speed was calculated by dividing the track length with the track duration.

#### Isolation of mouse bone marrow cells

Mice were anesthetized and sacrificed by cervical dislocation. Long bones (femurs, tibiae, humerus) were harvested into ice-cold sterile PBS. Bones were flushed with PBS + 2% fetal calf serum (FCS) using a 26-Gauge needle and the bone marrow suspension was further filtered through a 70µm cell strainer (Miltenyi Bioec GmbH, Bergisch Gladbach, Germany) and pelleted at 4℃, 300g for 5min. The supernatant was discarded and cells were resuspended and incubated in red blood cell lysis buffer for 5min. Lysis was terminated by adding 30ml PBS + 2mM Ethylenediaminetetraacetic acid (EDTA, Sigma-Aldrich, Saint Louis, MO, USA), followed by centrifugation 4℃ 300g for 5min. Cells were resuspended with PBS + 0.5% bovine serum albumin (BSA, Sigma-Aldrich, Saint Louis, MO, USA).

#### Megakaryocyte culture from mouse bone marrow

Bone marrow cells (see above) were cultured in DMEM medium containing 10% fetal bovine serum, 1% penicillin/streptomycin and 70 ng/µl thrombopoietin (TPO, ImmunoTools) for 5 days at 37℃ and 5% CO2. On day five a BSA step gradient was prepared by placing PBS containing 1.5% BSA on top of PBS with 3% BSA (PAA). Cells were loaded on top of the gradient, and MKs were settled to the bottom within 30 min at 1× gravity at room temperature. Mature MKs formed a pellet at the bottom of the tube.

#### In vitro co-culture of pDCs with MKs

pDC generation: bone marrow cells were isolated (see above) and cultured for 7 days in RPMI-1640 GlutaMAX-I (GIBCO) supplemented with 10% FCS (GIBCO), 1mM sodium pyruvate (GIBCO), 1% P/S (GIBCO), 1% NEAA (GE-Healthcare), 0.05 mM ß-mercaptoethanol MeEtOH (GIBCO) and recombinant 100ng/ml FLT3L (Biolegend). Cells were harvested by flushing petri dishes with cold PBS. Purity of pDCs was 70-75% as determined by FACS. *MK-iDTR* mice were injected with diphteria toxin (DT) to induce death of MKs. Control mice received PBS. After 6h of DT injection, mice were euthanized and femurs were flushed with DMEM medium containing 10% fetal bovine serum, 1% penicillin/streptomycin and 70 ng/ml thrombopoietin (TPO, ImmunoTools). MKs were isolated using a BSA gradient as described above. pDCs and MK (1:1) were incubated together for 18h at 37°C and 5% CO2. After incubation, the supernatant was harvested and analyzed for IFNα level (ELISA, see below).

#### MK colony forming unit (CFU) assay

CFU assays were performed using the MegaCult^TM^ kit (SteamCells Technologies) following the manufacturer’s protocol. In brief, femurs and tibias of *vWF-Cre+;IFNaR^-/-^* and *vWF-Cre-;IFNaR^-/-^*mice were flushed with Iscove’s MDM with 2% FBS to isolate bone marrow cells. Cells were washed in Iscove’s MDM (without FBS) prior to culture. 2,2 x 10^6^ cells were then resuspended in cold MegaCult™-C medium containing collagen, TPO 50 ng/ml and IFN alpha type 1 universal (5U, 10U, 100U, 500U or 1000U) (PBL-Biomedical Laboratories). The final cell suspension (1,5 ml) was loaded into 6-well plates and cultivated for 7 days at 37°C and 5% CO2. Following incubation, well plates were imaged with a stereo microscope Axio Zoom v16 from Carl Zeiss. MK-CFUs colonies were classified according to the manufacturer’s protocol (a minimum of 3 cells in close contact).

#### Flow cytometry

Cells were incubated with mouse CD16/CD32 (BD Pharmingen (Germany) (Fc block) before staining. The following antibodies were used to identify MKs: 1:100 anti-mouse CD41-FITC^+^ and anti-mouse CD42d-APC^+^ (BioLegend); and MKPs: anti-mouse CD41-FITC^+^, Pacific blue lineage negative (Ter-119-, CD3e-, CD45R-, CD11b-, Ly-6G-), anti-mouse CD105-PercCy7^-^, CD150-Brillint violet 510^+^ and anti-CD9-PercCy5.5^+^ (BioLegend) (all 1:100). We identified pDCs using the following antibodies: anti-mouse SiglecH-FITC^+^, CD11b-PE-Cy7-, B220-APC+ from BioLegend (San Diego, USA) (1:200). pDC activation: anti-mouse CD69-FITC, CD86-PE, CD11b-APC-Cy7, CD317-APC, SiglecH-PercCy5.5 antibodies all from BioLegend and Life/Dead marker (405 nm excitation; ThermoFisher). IFN expression by pDCs: Following staining with pDC surface markers (see above), cells were fixed with PFA and methanol and stained with monoclonal anti-mouse p-IRF7 antibody (cellsignaling, clone D6M2I) in Perm buffer III (BD) as previously described^31^. Before loading the samples, 10µg/ml Sytox Orange for the live/dead cell gating and counting beads (1,2,3 counting beads, ThermoFisher, Germany), were added to the cell suspension. Apoptosis was measured using Apotracker™ Green (BioLegend) according to manufacturer instructions. Measurements were performed on a Canto III cell analyzer (BD Biosciences, Germany) or Cytoflex-S (Beckman Coulter, Germany). FACS data was analyzed with FlowJo^TM^ 10.6.2. software.

#### EdU proliferation assay

Click-it® EdU Cell Proliferation Assay Kit (Thermo Fisher Scientific, Massachusetts, USA) was used to analyze the MKP proliferation. In vivo labeling of BM cells with 5-ethynyl-2’-deoxyuridine (EdU) was described previously^50^. Briefly*, vWF-eGFP* mice were intraperitoneally injected 0.5 mg EdU in DMSO. After 4h, mice were anesthetized and sacrificed by cervical dislocation and long bones (femurs and tibiae) were harvested. Bone marrow cells were prepared as described above. The detection of EdU was performed according to the manufacturer’s protocol. Briefly, cells were stained with surface marker antibodies (CD41, CD42) for 30 min RT in dark, followed by fixation for 15 min (4% PFA, provided in the kit) and permeabilization 15 min (saponin-based permeabilization and wash reagent, provided in the kit). Samples were washed with 1% BSA between each step. Samples were then incubated for 30min RT in dark in EdU reaction cocktail containing PBS, Copper protectant, Pacific Blue picolyl azide and reaction buffer additive, according to the manufacturer’s protocol. After samples were washed and analyzed by flow cytometry using a LSRFortessa cell analyzer (BD Biosciences, New Jersy, USA). vWF+ CD41+ CD42-cells were gated and EdU+ cells were measured within this population using FlowJo^TM^ 10.6.2. software (Ashland, USA).

#### ELISA

Serum TPO measurement: 1 ml anti-coagulated blood was collected intracardiacally and kept overnight at -20 ℃. The next day, the blood was centrifuged with 2000g for 20min and the supernatant (serum) was collected for TPO measurement using Quantikine Mouse Thrombopoietin ELISA Kit (R&D Systems, Minneapolis, USA) to measure serum TPO levels. Interferon alpha was measured by ELISA (Mouse IFN Alpha All Subtype ELISA Kit, High Sensitivity, PBL Assay Science). Blood was left at room temperature for 20 min and after centrifugation serum was frozen at -20°C until further analysis. To measure interferon levels in BM, one femur was flushed with 200 µl of PBS and cells were spun down at 300g. Supernatants were stored at -20°C until analysis.

#### RT-PCR of IFNaR

Megakaryocytes (MKs) and megakaryocyte progenitors (MKPs) from unfractionated murine bone marrow cell suspensions were directly sorted into RLT buffer (Qiagen, Hilden) containing 143 mM beta-mercaptoethanol (Sigma Aldrich) and total RNA was isolated using a RNeasy Micro Kit (Qiagen) including an on-spin column DNase I digest to remove remaining traces of genomic DNA. First-strand cDNA was synthesized from total RNA with the High Capacity cDNA Reverse Transcription kit (Applied Biosystems) using random primers in 20 µL reaction volumes. RT-PCR was performed using the SsoAdvanced Universal SYBR Green Supermix (BioRad) and the primers for murine IFNAR 1 and beta-Actin in a MyiQ™ Single-Color Real-Time PCR System (BioRad). Products of RT-PCR were separated by electrophoresis on a 2.5% agarose gel in 1x TBE buffer. Images were taken using a Gel iX Imager (Intas).

Primer:

Mm_Ifnar1 Fw TCTCTGTCATGGTCCTTTATGC Eurofins Mm_Ifnar1 Rev CTCAGCCGTCAGAAGTACAAG Eurofins Mm_Actb_1_SG primer assay (400 x 25 µL reactions), Cat#: QT00095242, Qiagen.

#### RNA-Seq analysis

For RNAseq analysis BM cells were isolated by flushing the long bones with FACS buffer (2mM EDTA, 1% FCS, PBS) and treated with Pharm LyseTM buffer (BD). Cells were enriched by magnetic removal of CD11b+ and CD19+ cells (EasySepTM, stem cell technologies). The negative fraction was stained for B220-BV421, SiglecH-PE, CD9-PerCP-Cy5.5, CD41-FITC, CD42-APC, cKit-APC-Cy7. 2000 cells were sorted into NEB-lysis buffer and processed for sequencing using the NEBNext® Single Cell/Low Input RNA Library Kit according to the manufacturer’s protocol (at IMGM, Martinsried, Germany). Libraries were pooled in equimolar amounts and sequenced on a NovaSeq6000 (Illumina) in a single-end 75nt run, yielding between 15 to 25 million reads per sample. Reads were mapped against GRCm38.p4 using CLC Genomics Workbench (Qiagen) with the following parameters: mismatch cost 2, insertion/deletion cost 3, length fraction 0.8, similarity fraction 0.8, global alignment “no”, strand specific “both”, maximum number of hits per read 5. CLC Genomics Workbench was also used for generation of gene expression matrices.

##### Gene set enrichment analysis (GSEA)

To prepare the data for GSEA, DESeq2 analysis was performed using Galaxy and default parameters^51, 52^. Genes were filtered for an expression of TPM>1 in any condition (42,868 genes) and then sorted according to the log2 fold change of the respective analysis. For further analysis, the tool GSEA (version 4.0.3) of UC San Diego and Broad Institute was used^53^ and ^54^, referring to their RNASeq manual pages for analysis. Gene sets of their Molecular Signatures Database (MSigDB) in the categories “Canonical Pathways” (C2) and “Gene Ontology” (C5) were chosen to contain Ifna1 gene (94 gene sets). Mouse gene symbols were mapped onto human gene symbol, gene set size was set to contain between 15-2000 genes.

##### Gene ontology analysis

Genes were filtered for log2 fold change greater or lower than 1 and submitted to the Database for Annotation, Visualization and Integrated Discovery (DAVID) version 6.8. Resulting gene ontology terms were filtered for a q-value <0.05.

##### Data visualization

ClustVis was used to generate the heatmap of genes expressed in megakaryocyte progenitors in Extended Data Figure 7 ^55^.

##### Data accessibility

RNA-Seq data will be accessible under a GEO database entry.

#### Statistics

GraphPad Prism software (9.1.2, San Diego, USA) was used for all statistical analysis. All data were assumed to have Gaussian distribution, unless specified. Before doing statistical analysis, the data were confirmed to have equal variance using F-test, and Student’s unpaired t-test was used for the comparison of two groups; otherwise, unpaired t-test with Welch’s correction was used when variances are significantly different. For comparison of multiple groups, one-way ANOVA was used. Error bars indicate standard deviation (SD) or standard error of the mean (SEM). P values less than 0.05 were considered significant.

## Data availability

Data that supports the findings of this study are available within the article and its Supplementary Information. Any additional information and related data are available upon reasonable request.

## Acknowledgements

We thank Sebastian Helmer, Nicole Blount, Elisabeth Raatz, Zeljka Sisic, Anna Titova for excellent technical assistance. This work was supported by the German Research Foundation (SFB 1123 [S.M. project B06]; SFB 914 [S.M. projects B02 and Z01, H.I.-A. project Z01, S.S. project A06, F.G. Gerok position, K.S. project B02, C.Schu. project A10, B.W. project A02, C.Schei. project B09]; SFB 1054 (T.B.); FOR2033 [F.G., R.A.J.O., S.M.]; by the DZHK (German Center for Cardiovascular Research) (MHA 1.4VD [S.M.], Postdoc Start-up Grant, 81X3600213 [F.G.]); by FöFoLe project 947 (F.G.), the Friedrich-Baur-Stiftung project 41/16 (F.G.), LMUexcellence NFF (F.G.). P.B. is supported by the German Research Foundation (Project-IDs 322900939 (SFB TRR 219), 454024652, 432698239, and 445703531 and 445703531 (CRU 5011), European Research Council (ERC) Consolidator Grant AIM.imaging.CKD (No. 101001791), the German Registry of COVID-19 autopsies (www.DeRegCOVID.ukaachen.de) funded by the Federal Ministry of Health (ZMVI1-2520COR201) and the Federal Ministry of Education and Research (STOP-FSGS-01GM1901A and DEFEAT PANDEMIcs, No. 01KX2021). S.v.S. is supported by the START-Program of the Faculty of Medicine of the RWTH Aachen University (AZ 125/17). This project has received funding from the European Research Council (ERC) under the European Union’s Horizon 2020 research and innovation programme (grant agreement No. 833440 to S.M.). F.G. received funding from the European Union’s Horizon 2020 research and innovation programme under the Marie Skłodowska-Curie grant agreement no. 747687.

## Author contributions

Initiation, F.G.; Conceptualization, F.G. and S.M. with input from H.I-A., S.S.; Methodology, H.I-A., S.S., W.F., F.G., C. Schei., A.D., V.F., M.I., C.Gi..; Investigation, H.I.-A., S.S., W.F., C.Gu., J.W., Z.Z., D.v.d.H., A.D., T.S., V.F., M.L., C. Schu., C.Gi., M.R., Resources, K.S., M.C., T.B., B.W., S.E., W.C.A., T.P., M.S., C.N., M.I., R.A.J.O., S.M.; Formal Analysis, H.I.-A., S.S., W.F., A.D., C.Gi.; Writing – Original Draft, F.G.; Writing – Editing, F.G., S.M., H.I.-A., S.S. with input from all authors; Visualization, H.I-A., S.S., W.F., F.G., A.D., D.v.d.H.; Supervision, F.G., H.I.-A., S.M.; Project Administration, F.G; Funding Acquisition, F.G. and S.M.

## Material & Correspondence

Further information and requests for resources and reagents should be directed to and will be fulfilled by the Lead Contact, Florian Gaertner.

## Supplementary figures with legends

**Extended Data Figure 1.**
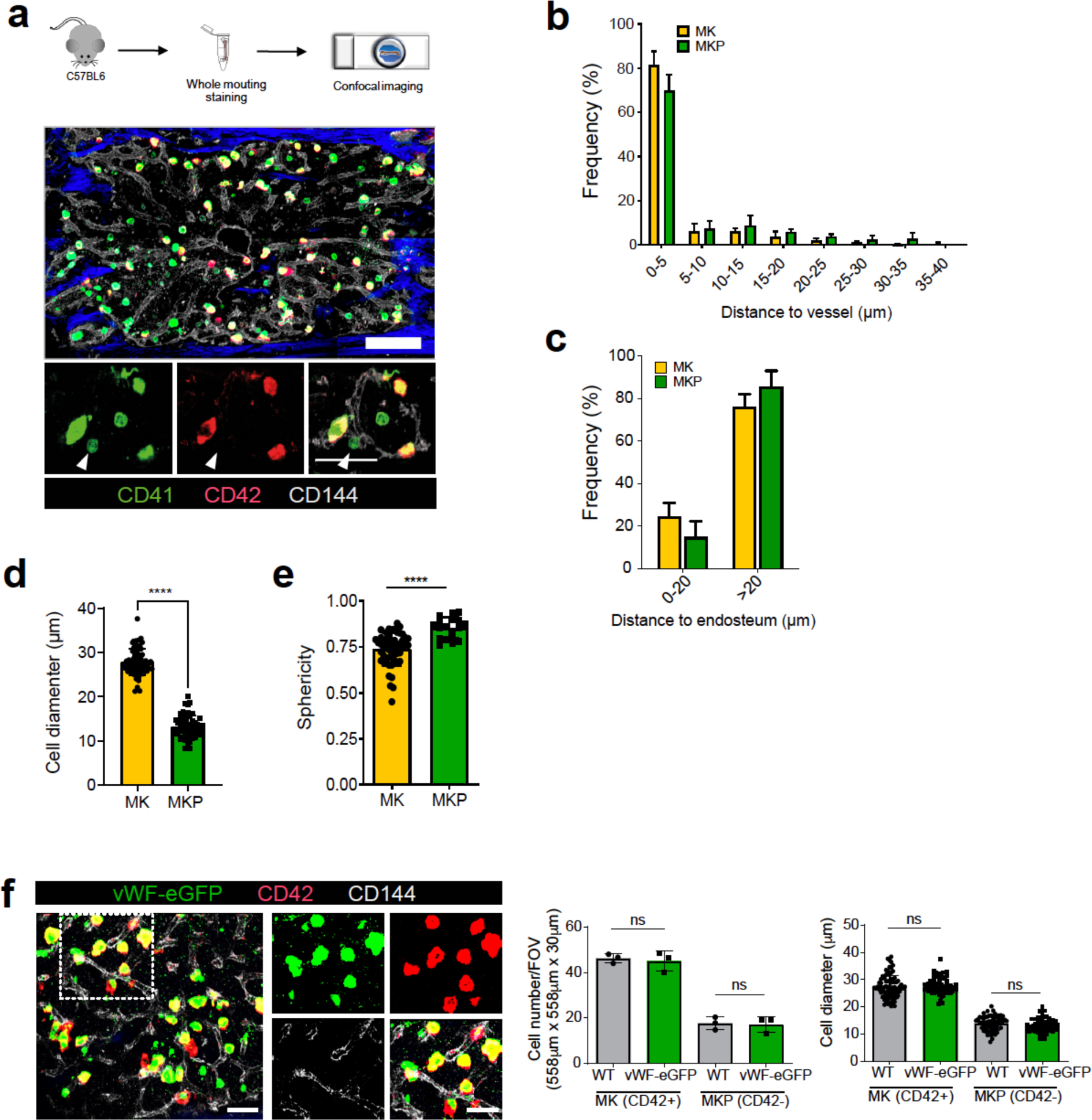
MK and MKP distribution in the bone marrow niche. **a**, Representative whole-mount immunostaining of megakaryocyte progenitors and mature megakaryocytes in murine sternum bone. MKPs (green): CD41^+^/CD42^-^; MKs (yellow): CD41^+^/CD42^-^; blood vessels (grey): CD144^+^; bone (blue): second harmonic generation. Arrowhead indicates MKP. Scale bars = 200 µm (upper); 100 µm (lower). **b**, **c**, Histograms showing BM distribution of MKs and MKPs relative to their distance to vessels and endosteum. n=3; Mean±SEM. **e**, Cell diameter and sphericitity; for cell diameter MK n= 68 cells and MKP n=55 cells; for sphericity MK n= 53 cells and MKP n= 17 cells; pooled from 7 mice; ****: p<0.0001; unpaired t-test with Welch’s correction; Mean±SEM. **f**, Representative whole-mount immunostaining of MKs/MKPs in vWF^eGFP/+^ mice. MKs/MKPs of vWF^eGFP/+^ mice show no significant difference in cell number and size compared to C57Bl/6J mice. n= 3 mice; unpaired t-test; ns= no significance; Mean±SEM. Scale bar=30µm. **g**, vWF-eGFP does not co-localize with erythrocytes (Ter-119), granulocytes (CD11b, Ly-6G) and lymphocytes (CD3e, CD45R) in BM. Scale bar=100 µm. **h**, Platelet counts of vWF^eGFP/+^ compared to C57Bl/6J; WT group n=14, vWF-eGFP group n=7. Unpaired t-test; Mean±SEM. n.s.= no significance (p=0.1369).

**Extended Data Figure 2.**
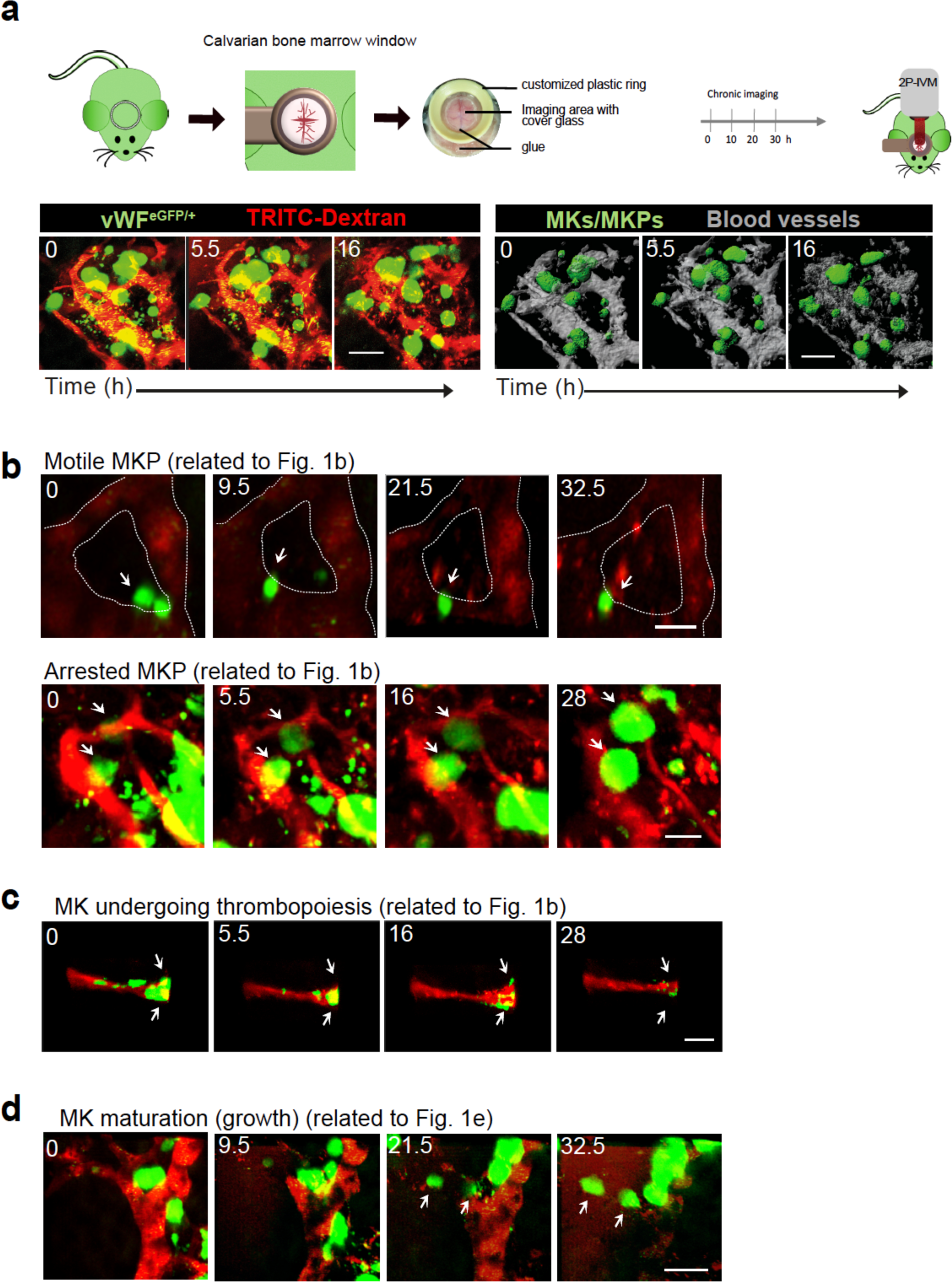
Megakaryocytic lineage tracing by chronic time-lapse 2P-IVM. **a,** Schematic showing experimental setup of chronic 2-photon intravital microscopy of mouse calvarian bone marrow. vWF^eGFP/+^ mice were i.v. injected with TRITC-dextran to track the megakaryocytic lineage (green) and blood vessels (red) respectively (left time series). Right: 3D rendering of z-stacks from the same 2P-IVM time series. MK/MKPs are depicted in green and blood vessels are false colored in grey (right). **b**-**d**, Raw data corresponding to 3D-rendered images shown in Figure 1. Scale bars= 50 µm.

**Extended Data Figure 3.**
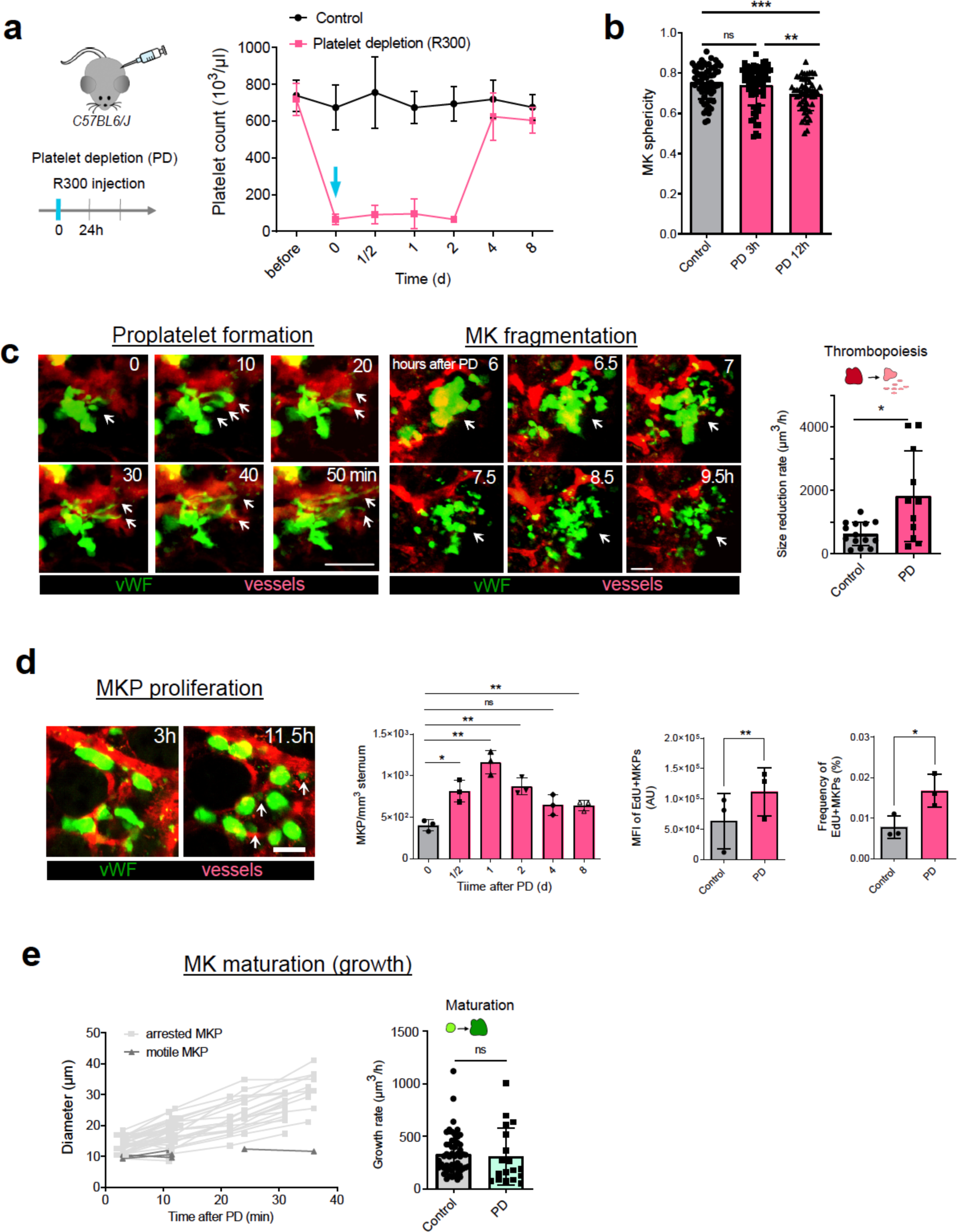
Spatio-temporal dynamics of the megakaryocytic lineage in response to immune-mediated thrombocytopenia. **a,** Schematic showing experimental setup of PD. Peripheral blood platelet counts monitored at indicated timepoints after a single injection (i.p.) of R300 or isotype control; n≥3 mice; Mean±SEM. **b**, Morphometric analysis of MKs shows decreased sphericity in response to PD corresponding to an increase of cellular protrusions (proplatelets); Control n= 63 cells, PD 3h n= 65 cells and PD 12h n=51 cells, pooled from 4 mice; unpaired t-test ns: p=0.3557, **: p= 0.0097, ***: p=0,0002. Mean±SEM. **c,** Representative 2P-IVM time series of proplatelet formation and MK fragmentation. Single cell tracking of MK volumes over time reveals a significant faster decrease of volume following PD. Control n=14 and PD n=11 cells, pooled from 3 mice; unpaired t-test; ns: p=0.0206; Mean±SD. **d,** Left: Small (<15μm) vWF-eGFP+ cells appear 12h after PD (2P-IVM). Scale bar=50 µm. Middle: PD triggers an instantaneous proliferation of MKPs peaking 1d following platelet depletion; MKPs (CD41+/CD42-) were counted in whole mount BMs. n= 3 mice; unpaired t-test, ns: p= 0.0531, *: p=0.0184, **: p=0.0042, **: p=0.0045 and **: p=0.0100; Mean±SD. Right: Proliferation of MKPs (vWF-eGFP+/CD41+/CD42-) was measured after in vivo labeling of BM cells with 5-ethynyl-2’-deoxyuridine (EdU) using FACS. Mean fluorescent intensity of EdU and Frequency of EdU-pve cells significantly increases after PD (12h); n=3 mice; paired t-test; *: p=0.0046; **: p=0.0115; error bar=SD; Mean±SD. **e,** Left: MKPs lodged within the perivascular niche of the BM grow in volume. Volume increase of single cells was tracked over time. Arrested MKP n=27 cells and mobile MKP n=6 cells. Notably, the speed of cell growth after PD did not significantly differ from steady state control; Control n=54 cells and PD= 18 cells, pooled from 3 mice, unpaired t-test Welch’s correction ns:p= 0.7503; Mean±SD.

**Extended Data Figure 4.**
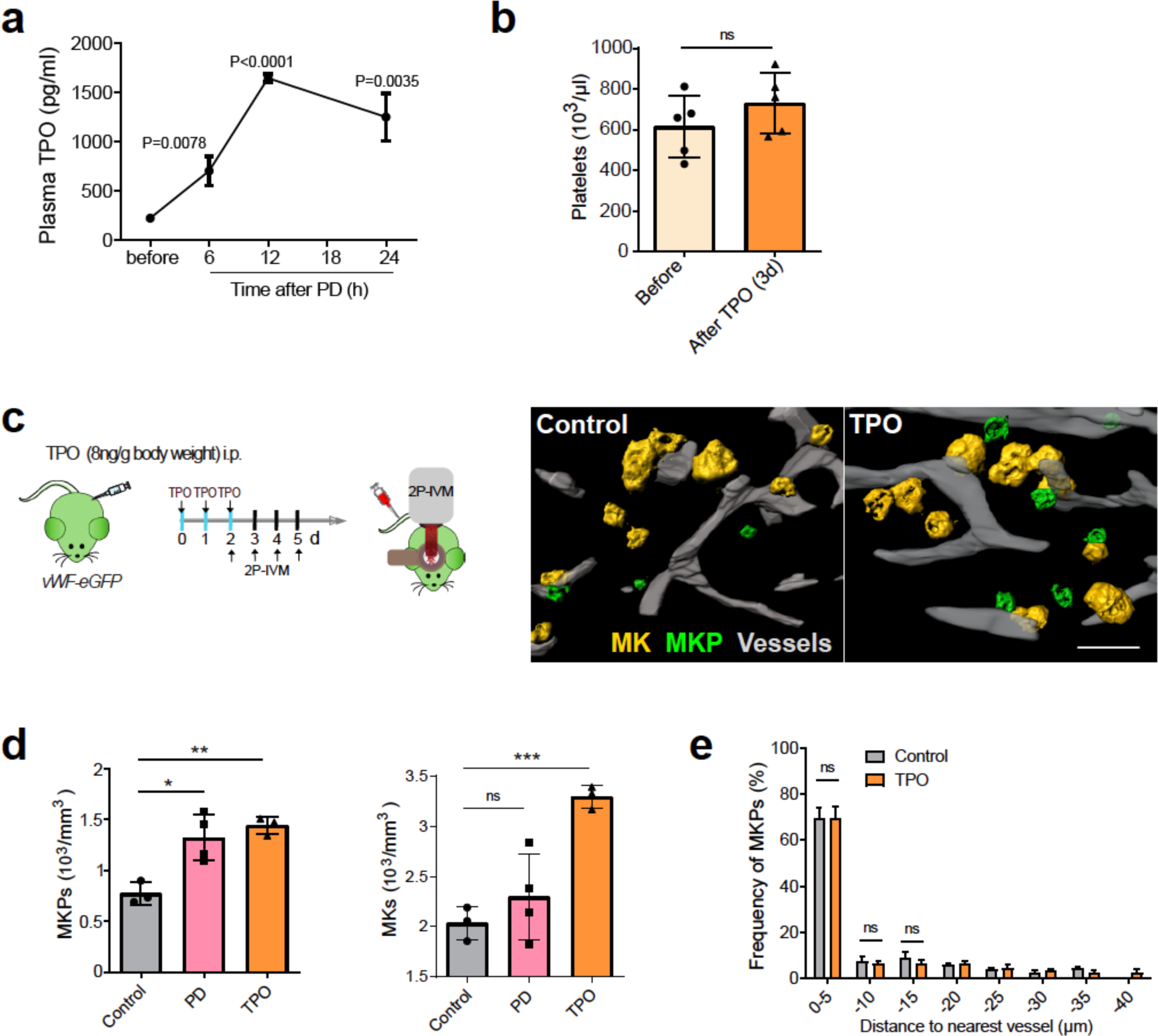
TPO triggers global megakaryopoiesis without preferential localization to the perivascular niche. **a,** Plasma TPO levels increase in response to PD, reaching the highest levels 12h after platelet depletion (ELISA); n=4 mice per group, unpaired t-test with Welch’s; Mean±SEM. **b,** Thrombopoiesis was unaffected by TPO treatment (8ng/g body weight on 3 consecutive days, i.p.) as indicated by unaffected platelet counts (hemocytometer); n=5 mice, unpaired t-test Welch’s correction ns: p=0.267; Mean±SD. **c**, Representative micrographs of chronic 2P-IVM show increased numbers of MKPs (green: vWF-eGFP^+^/<15 μm) and MKs (green: vWF-eGFP^+^/>15 μm); scale bar=50 µm. **d**, TPO-treatment increased megakaryopoiesis (MKP numbers) to an extent similar to platelet depletion (PD). TPO-treatment leads to an accumulation of mature MKs in the BM while MK numbers in the BM remained unaffected after PD due to increased MK consumption (BM whole-mount immunostainings); control n= 3 mice, PD n=4 mice and TPO n=3 mice, unpaired t-test; MKPs, *: p=0.0106, **: p= 0.0015; MK, ns=0.3234, ***: p=0.0008; Mean±SD. **e**, Increased megakaryopoiesis in response to TPO followed an entirely different pattern compared to PD. During PD we found a local increase in megakaryopoiesis confined to the perisinusoidal compartment (**see** Fig. 1g). In contrast, TPO treatment boosts megakaryopoiesis across the entire BM compartment and disrupts preferential perivascular MK replenishment from MKPs. Consequently, distribution of MKPs relative to the perivascular niche was unaffected by TPO treatment as analyzed in BM whole-mount immunostainings. The distances were binned into 5 µm intervals. n=4 mice per group. Unpaired t-test; ns: p=0.7, ns: p=0.9. Mean±SD.

**Extended Data Figure 5.**
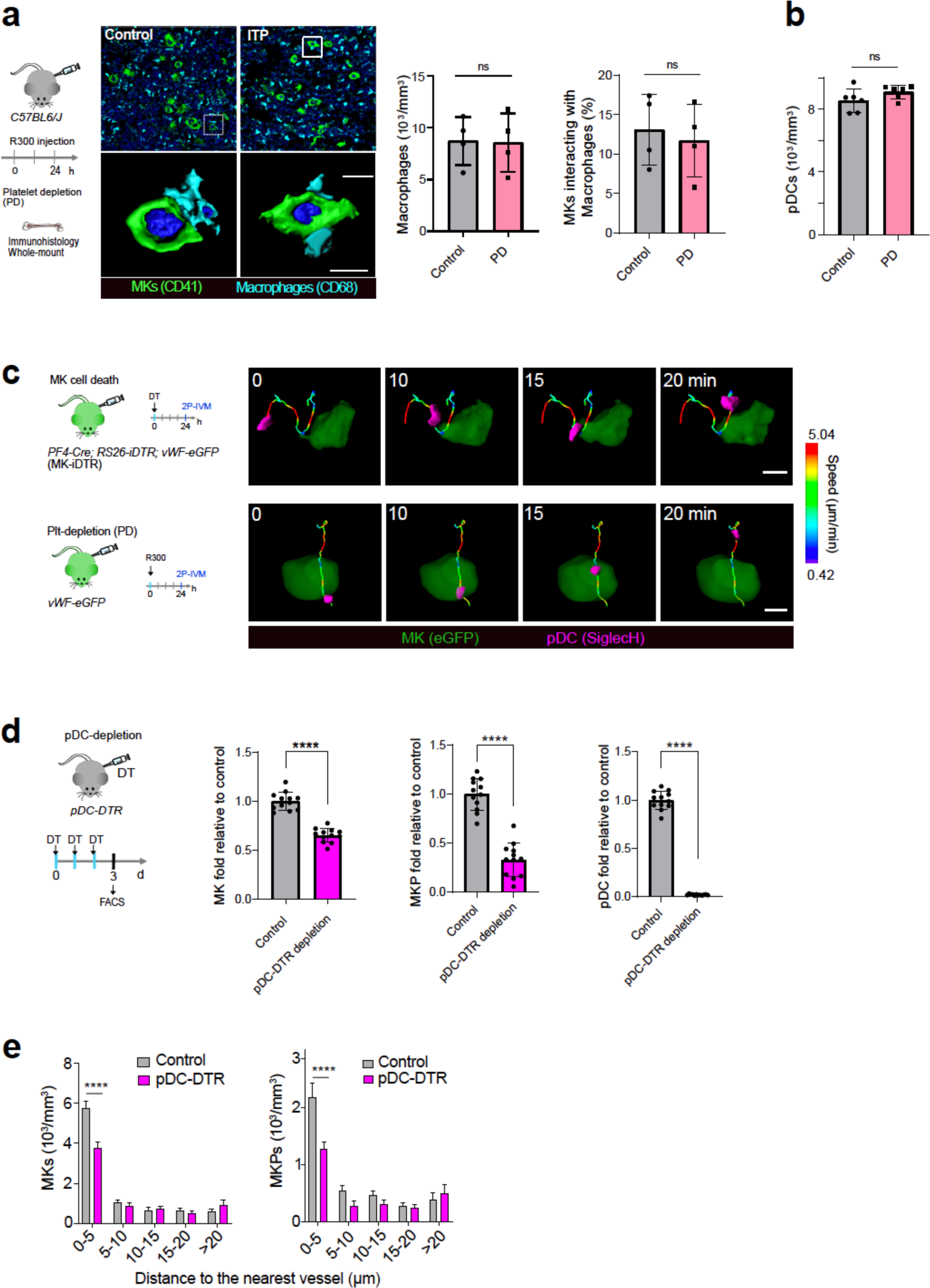
pDCs are BM niche cells that regulate megakaryopoiesis. **a**, MK-Macrophage interactions were quantified by whole-mount BM immunohistology; MK: CD41+; Macrophages: CD68+. Scale bar=10 µm. PD did not increase MK-Macrophage interactions; n= 4 mice; unpaired t-test Welch’s correction ns: p=0.9299 and ns: p=0.6882; Mean±SD. **b**, PD did not increase the number of bone marrow pDCs, as counted in whole-mount BM immunohistology (pDCs: BST2+); n= 6 mice; unpaired t-test Welch’s correction ns: p=0.1537. Mean±SD. **c**, Left: Schematic showing experimental setup of 2P-IVM of pDC and MK interactions. Right. Representative time series of pDCs interacting with MKs. Color-code: speed of migratory pDCs. Scale bars=10 µm**. d**, Impaired megakaryopoiesis following pDC-depletion in *BDCA2-DTR (pDC-DTR)* mice (also see Methods). Cell numbers were quantified by FACS (see Extended Fig. 6 for gating strategy); n=12 mice; unpaired t-test Welch’s correction, ****: p<0.0001; Mean±SD. **e,** pDC-depletion disrupts the megakaryocytic niche and alters the distribution of MKPs and MKs within the BM. BM whole-mounts were stained for MKs (CD41+CD42+) and MKP (CD41+CD42-) and positioning was quantified in relation to blood vessels (CD144+). Numbers of MKs and MKPs in close contact to blood vessels significantly decreased following pDC-depletion; n=12 mice; unpaired t-test Welch’s correction, ****: p<0.0001. Mean±SD.

**Extended Data Figure 6.**
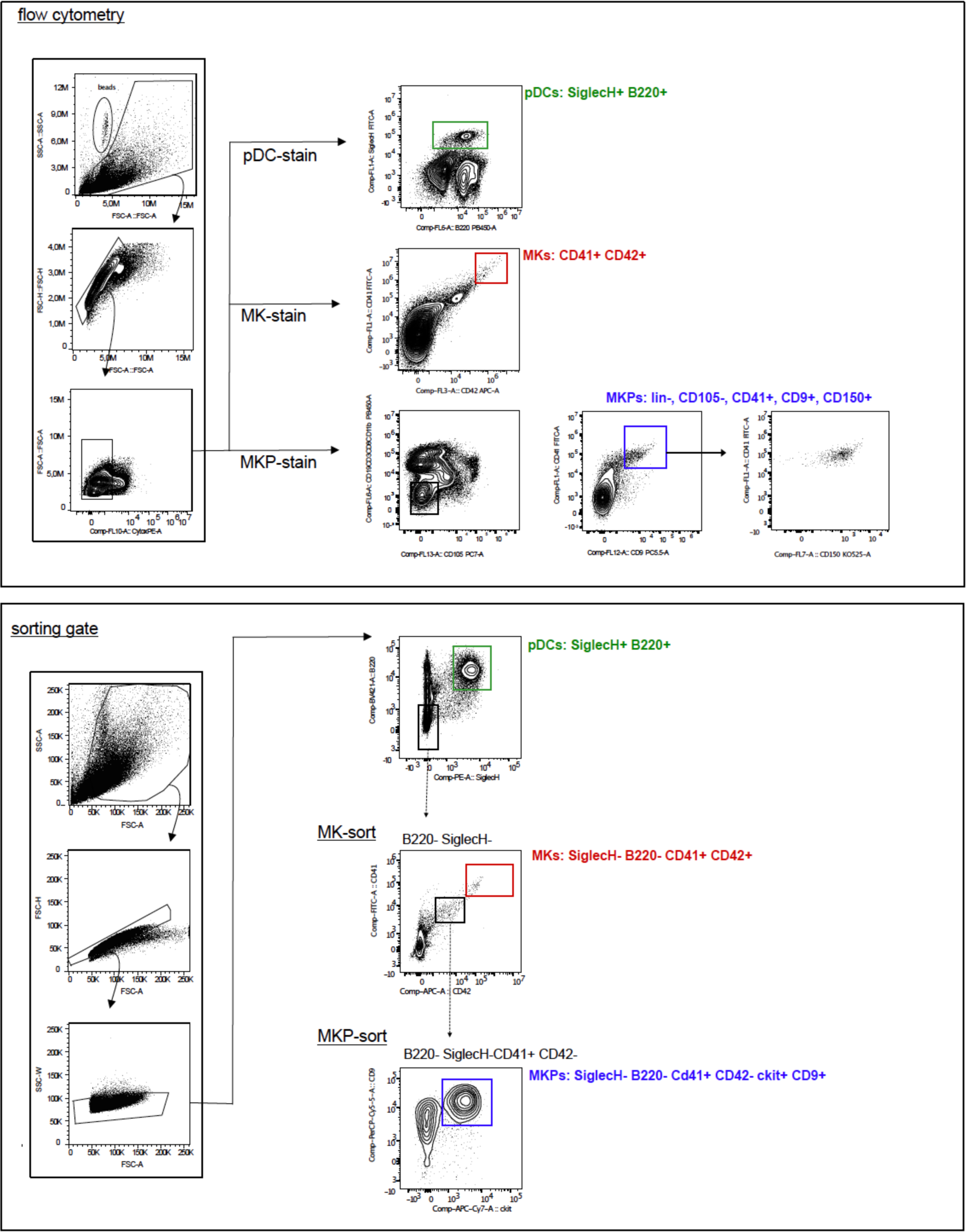
Gating strategy for pDC-, MK- and MKp-analysis. Upper panel shows the identification of pDCs, MKs and MKps by staining of living cells for SiglecH, B220, CD41, CD42 and lineage negative cells for CD41 and CD9. Counting beads were used during FACS analysis. Lower panel demonstrates the sorting strategy of SiglecH-, B220-cells that are CD42 neg CD41+ and CD9 and ckit+.

**Extended Data Figure 7.**
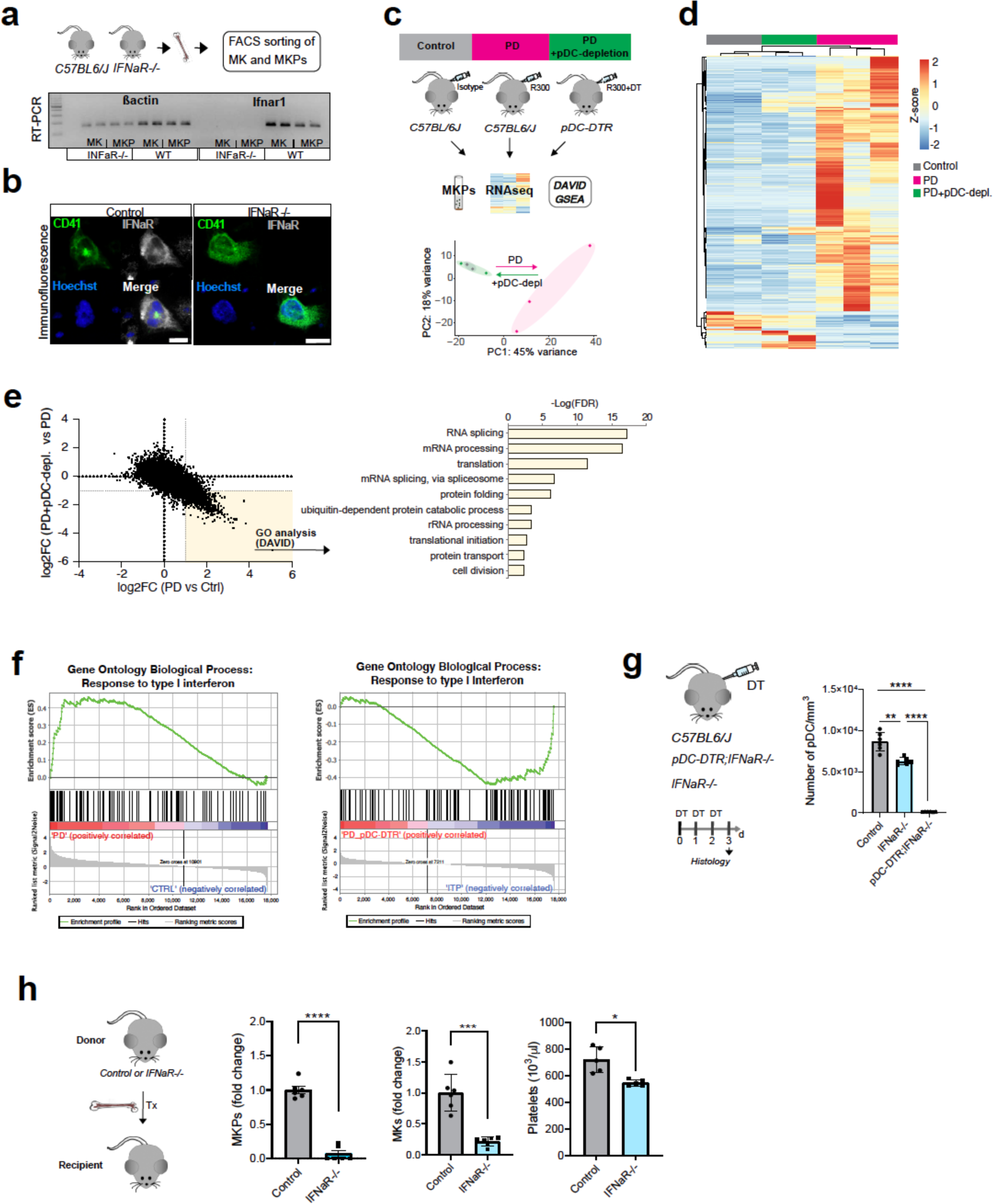
pDC-derived IFNα fuels megakaryopoiesis. **a**, Expression (mRNA) of *IFNaR* was analyzed in sorted MKs and MKPs by RT-PCR. MKPs and MKs from *IFNaR*-/- mice served as negative control; *ß-actin* (housekeeping gene) **b**, Immunofluorescence staining confirmed expression of IFNaR in MKPs and MKs. *IFNaR-/-* (negative control). Scale bar=10 µm. **c**, Upper: Schematic of experimental design. Lower: Principal component analysis (PCA) of MKP RNA-Seq data. MKPs exhibit a strong variance shift following PD (grey to magenta). This variance is abrogated when animals were additionally depleted for pDCs (magenta to green). **d**, Expression heatmap of MKP RNA-Seq data. Heatmap shows MKP genes de-regulated (log2FC <1 or >1 with FDR<0.05) in either PD versus control or PD vs. platelet depletion with additional pDC depletion. Heatmap was generated using non-hierarchical clustering on rows and columns using ClustVis R package. **e,** Cell proliferation-associated genes are inversely regulated between platelet depletion and additional pDC depletion. Scatter plot shows de-regulated genes (log2FC) in PD vs. Control and PD + pDC-depletion vs. PD plotted against each other. Common genes were submitted to gene ontology analysis (DAVID), all significant terms for bioprocesses (FDR<0.05) are shown. Of note, DAVID revealed upregulated genes associated with terms for activated transcription, translation and cell proliferation in general (Top 10 terms are shown), in accordance to MKP proliferation observed after PD. **f,** Gene set enrichment analysis was performed on RNA-Seq data of MKPs after PD compared to control (left) as well as after PD with additional pDC depletion (right), respectively. **g,** Decreased MKP and MK numbers in *IFNaR-/-* mice. Notably, depletion of pDCs in *pDC-DTR;IFNaR-/-* mice had no additive effect. Unpaired t-test ns: p=0.2589. pDCs, MKPs, MKs were counted immunofluorescent labelling of BM sections); n= 6 mice; unpaired t-test; *: p=0.0339, **: p=0.0027 and 0.0012, ***: p=0.0009, ****: p<0.0001. Mean±SD. **h,** Decreased MKP and MK numbers in BM chimeric *IFNaR-/-* mice. Irradiated WT mice received BM from *IFNaR-/-* or *IFNaR+/+ (Control)* donors, respectively. Mice were subjected to analysis 8 weeks after transplantation.

**Extended Data Figure 8.**
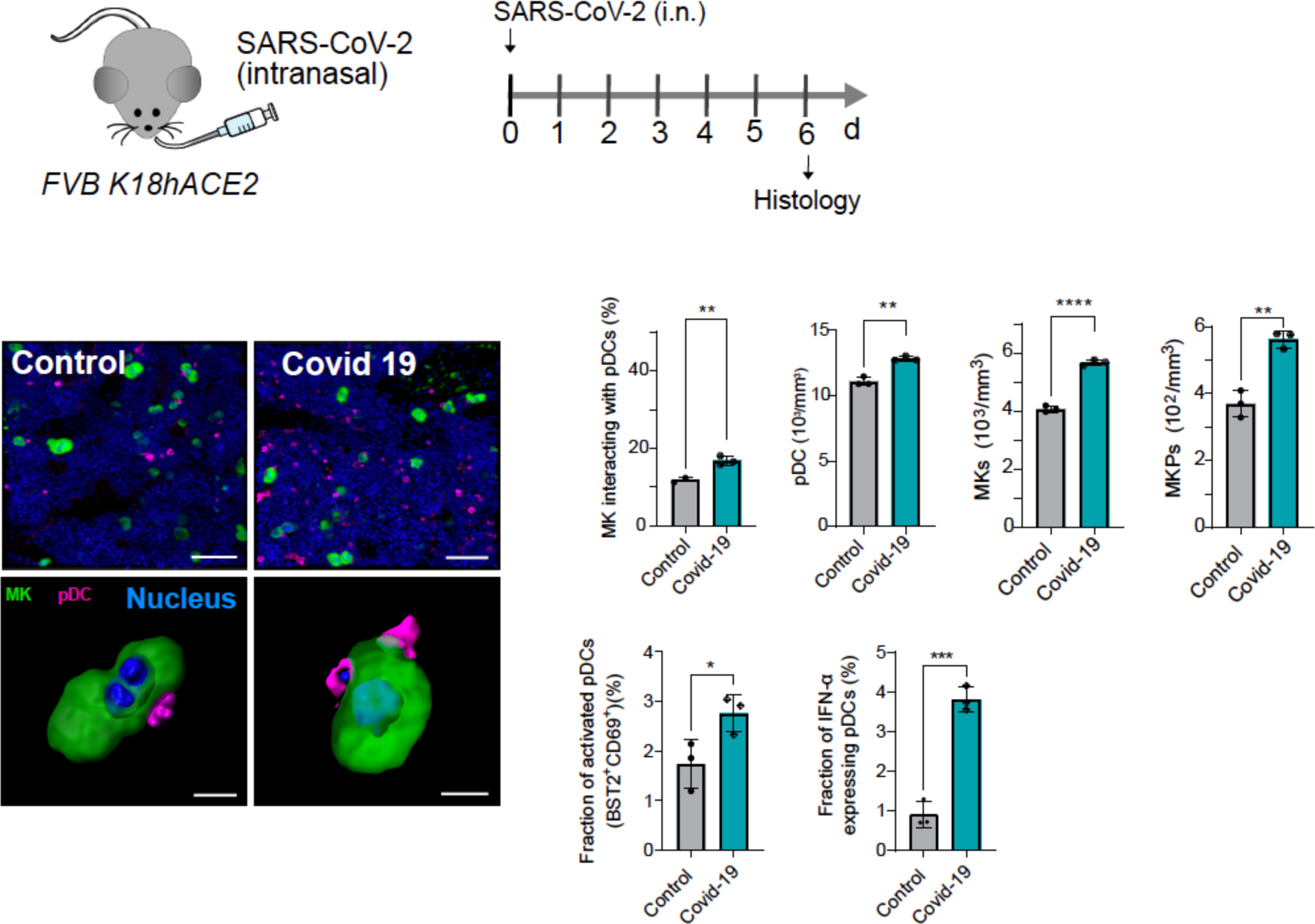
Augmented megakaryopoiesis in a humanized mouse model of SARS-CoV2 infection. BM from *FVB;K18hACE2* mice infected with SARS CoV-2 (10^5^ TCID_50_ SARSCoV2/mouse in 25μl intranasally or untreated control mice were analyzed by immunohistology. MKs (green; CD41;,>15μm); MKPs (green; CD41; <15μm); pDCs (magenta; BST2). MK/pDc interaction: n= 3 mice, **: p=0.0081, pDC number: **: p=0.0037, MK number: ****: p<0.0001, MKp **: p=0.0039.

### Supplementary video legends

**Supplementary video 1** MKs and MKPs reside along BM sinusoids. 2PM of sternal whole-mount (2D slices and 3D rendered stack are shown and animated). MKs (orange; CD42+/CD41+); MKPs (green; CD42-/CD41+); sinusoids (grey; CD144+); bone (blue; SHG).

**Supplementary video 2** Thrombosis and megakaryopoiesis are synchronized processes whin the megakaryocytic bone marrow niche. Chronic 2P-IVM of the cavarial bone marrow reveals mature MKs (color-coded in magenta) that disappear from the niche and small MKPs (color-coded in green) that appear and grow in size. The total number of megakaryocytic (vWF-eGFP+ cells) remains constant over time. Movie shows animation of 3D rendered data. SD-stacks were recorded at the four indicated timepoints. MKs are labeled with vWF-eGFP; blood vessels are visualized by TRITC-dextran. Also see Fig. 1e and Extended data Fig. 2d.

**Supplementary video 3** Migrating pDCs monitor megakaryocytes in the bone marrow. 2P-IVM movies of pDCs (magenta; anti-BST2-PE) interacting with MKs (green; vWF-eGFP^+^) in the bone marrow. 3D rendered data is shown. Representative pDC interactions with control MKs (untreated); with dying MKs (MK-iDTR-mice); and MKs during accelerated platelet demand (platelet depletion) are shown. In all examples, pDCs engage in close contact with MKs. pDC protrusions poke into MKs and sometimes pDCs even migrate through the MK cell body (emperipolesis). Also see Figure 2D and Extended Data Fig. 5c.

**Supplementary Table 1:**
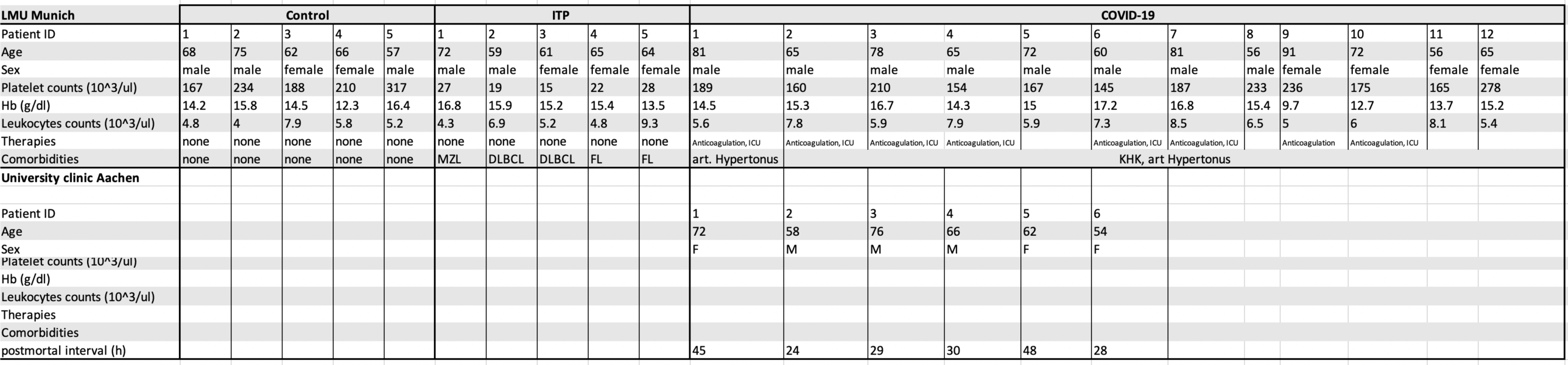
Patient characteristics.

## References

1. van der Meijden, P. E. J. & Heemskerk, J. W. M. Platelet biology and functions: new concepts and clinical perspectives. Nat Rev Cardiol 16, 166–179, doi:10.1038/s41569-018-0110-0 (2019).

2. Engelmann, B. & Massberg, S. Thrombosis as an intravascular effector of innate immunity. Nat Rev Immunol 13, 34–45, doi:10.1038/nri3345 (2013).

3. Machlus, K. R. & Italiano, J. E., Jr. The incredible journey: From megakaryocyte development to platelet formation. J Cell Biol 201, 785–796, doi:10.1083/jcb.201304054 (2013).

4. Junt, T. et al. Dynamic Visualization of Thrombopoiesis Within Bone Marrow. Science 317, 1767, doi:10.1126/science.1146304 (2007).

5. Reizis, B. Plasmacytoid Dendritic Cells: Development, Regulation, and Function. Immunity 50, 37–50, doi:10.1016/j.immuni.2018.12.027 (2019).

6. Gu, S. X. et al. Thrombocytopathy and endotheliopathy: crucial contributors to COVID-19 thromboinflammation. Nat Rev Cardiol 18, 194–209, doi:10.1038/s41569-020-00469-1 (2021).

7. Kunisaki, Y. et al. Arteriolar niches maintain haematopoietic stem cell quiescence. Nature 502, 637–643, doi:10.1038/nature12612 (2013).

8. Sanjuan-Pla, A. et al. Platelet-biased stem cells reside at the apex of the haematopoietic stem-cell hierarchy. Nature 502, 232–236, doi:10.1038/nature12495 (2013).

9. Stegner, D. et al. Thrombopoiesis is spatially regulated by the bone marrow vasculature. Nat Commun 8, 127, doi:10.1038/s41467-017-00201-7 (2017).

10. Eto, K. & Kunishima, S. Linkage between the mechanisms of thrombocytopenia and thrombopoiesis. Blood 127, 1234-1241, doi:10.1182/blood-2015-07-607903 (2016).

11. Pinho, S. et al. Lineage-Biased Hematopoietic Stem Cells Are Regulated by Distinct Niches. Dev Cell 44, 634–641 e634, doi:10.1016/j.devcel.2018.01.016 (2018).

12. McArthur, K., Chappaz, S. & Kile, B. T. Apoptosis in megakaryocytes and platelets: the life and death of a lineage. Blood 131, 605–610, doi:10.1182/blood-2017-11-742684 (2018).

13. Kaur, S. et al. Role of bone marrow macrophages in controlling homeostasis and repair in bone and bone marrow niches. Semin Cell Dev Biol 61, 12-21, doi:10.1016/j.semcdb.2016.08.009 (2017).

14. Gilliet, M., Cao, W. & Liu, Y. J. Plasmacytoid dendritic cells: sensing nucleic acids in viral infection and autoimmune diseases. Nat Rev Immunol 8, 594-606, doi:10.1038/nri2358 (2008).

15. Iparraguirre, A. et al. Two distinct activation states of plasmacytoid dendritic cells induced by influenza virus and CpG 1826 oligonucleotide. Journal of Leukocyte Biology 83, 610–620, doi:https://doi.org/10.1189/jlb.0807511 (2008).

16. Swiecki, M. & Colonna, M. The multifaceted biology of plasmacytoid dendritic cells. Nat Rev Immunol 15, 471–485, doi:10.1038/nri3865 (2015).

17. Cunin, P. et al. Megakaryocyte emperipolesis mediates membrane transfer from intracytoplasmic neutrophils to platelets. Elife 8, doi:10.7554/eLife.44031 (2019).

18. Bruns, I. et al. Megakaryocytes regulate hematopoietic stem cell quiescence through CXCL4 secretion. Nat Med 20, 1315-1320, doi:10.1038/nm.3707 (2014).

19. Zhao, M. et al. Megakaryocytes maintain homeostatic quiescence and promote post-injury regeneration of hematopoietic stem cells. Nat Med 20, 1321-1326, doi:10.1038/nm.3706 (2014).

20. Swiecki, M., Gilfillan, S., Vermi, W., Wang, Y. & Colonna, M. Plasmacytoid dendritic cell ablation impacts early interferon responses and antiviral NK and CD8(+) T cell accrual. Immunity 33, 955-966, doi:10.1016/j.immuni.2010.11.020 (2010).

21. Swiecki, M. & Colonna, M. Unraveling the functions of plasmacytoid dendritic cells during viral infections, autoimmunity, and tolerance. Immunol Rev 234, 142–162, doi:10.1111/j.0105-2896.2009.00881.x (2010).

22. Iannacone, M. et al. Subcapsular sinus macrophages prevent CNS invasion on peripheral infection with a neurotropic virus. Nature 465, 1079-1083, doi:10.1038/nature09118 (2010).

23. Cervantes-Barragan, L. et al. Control of coronavirus infection through plasmacytoid dendritic-cell-derived type I interferon. Blood 109, 1131–1137, doi:10.1182/blood-2006-05-023770 (2007).

24. Sa Ribero, M., Jouvenet, N., Dreux, M. & Nisole, S. Interplay between SARS-CoV-2 and the type I interferon response. PLoS Pathog 16, e1008737, doi:10.1371/journal.ppat.1008737 (2020).

25. Onodi, F. et al. SARS-CoV-2 induces human plasmacytoid predendritic cell diversification via UNC93B and IRAK4. J Exp Med 218, doi:10.1084/jem.20201387 (2021).

26. Laing, A. G. et al. A dynamic COVID-19 immune signature includes associations with poor prognosis. Nat Med 26, 1623–1635, doi:10.1038/s41591-020-1038-6 (2020).

27. Roncati, L. et al. A proof of evidence supporting abnormal immunothrombosis in severe COVID-19: naked megakaryocyte nuclei increase in the bone marrow and lungs of critically ill patients. Platelets 31, 1085-1089, doi:10.1080/09537104.2020.1810224 (2020).

28. Blanco-Melo, D. et al. Imbalanced Host Response to SARS-CoV-2 Drives Development of COVID-19. Cell 181, 1036–1045 e1039, doi:10.1016/j.cell.2020.04.026 (2020).

29. Bernardes, J. P. et al. Longitudinal Multi-omics Analyses Identify Responses of Megakaryocytes, Erythroid Cells, and Plasmablasts as Hallmarks of Severe COVID-19. Immunity 53, 1296–1314 e1299, doi:10.1016/j.immuni.2020.11.017 (2020).

30. Mesev, E. V., LeDesma, R. A. & Ploss, A. Decoding type I and III interferon signalling during viral infection. Nat Microbiol 4, 914–924, doi:10.1038/s41564-019-0421-x (2019).

31. Stutte, S. et al. Type I interferon mediated induction of somatostatin leads to suppression of ghrelin and appetite thereby promoting viral immunity in mice. Brain Behav Immun 95, 429-443, doi:10.1016/j.bbi.2021.04.018 (2021).

32. Gough, D. J., Messina, N. L., Clarke, C. J., Johnstone, R. W. & Levy, D. E. Constitutive type I interferon modulates homeostatic balance through tonic signaling. Immunity 36, 166–174, doi:10.1016/j.immuni.2012.01.011 (2012).

33. Essers, M. A. et al. IFNalpha activates dormant haematopoietic stem cells in vivo. Nature 458, 904–908, doi:10.1038/nature07815 (2009).

34. Sato, T. et al. Interferon regulatory factor-2 protects quiescent hematopoietic stem cells from type I interferon-dependent exhaustion. Nat Med 15, 696–700, doi:10.1038/nm.1973 (2009).

35. Iannacone, M. et al. Platelets prevent IFN-alpha/beta-induced lethal hemorrhage promoting CTL-dependent clearance of lymphocytic choriomeningitis virus. Proc Natl Acad Sci U S A 105, 629–634, doi:10.1073/pnas.0711200105 (2008).

36. Woo, A. J. et al. Developmental differences in IFN signaling affect GATA1s-induced megakaryocyte hyperproliferation. J Clin Invest, doi:10.1172/JCI40609 (2013).

37. Haas, S. et al. Inflammation-Induced Emergency Megakaryopoiesis Driven by Hematopoietic Stem Cell-like Megakaryocyte Progenitors. Cell Stem Cell 17, 422–434, doi:10.1016/j.stem.2015.07.007 (2015).

38. Martin, T. G. & Shuman, M. A. Interferon-induced thrombocytopenia: Is it time for thrombopoietin? Hepatology 28, 1430-1432, doi:https://doi.org/10.1002/hep.510280536 (1998).

39. Yamane, A. et al. Interferon-alpha 2b-induced thrombocytopenia is caused by inhibition of platelet production but not proliferation and endomitosis in human megakaryocytes. Blood 112, 542–550, doi:10.1182/blood-2007-12-125906 (2008).

40. Pinho, S. & Frenette, P. S. Haematopoietic stem cell activity and interactions with the niche. Nat Rev Mol Cell Biol 20, 303–320, doi:10.1038/s41580-019-0103-9 (2019).

41. Gaertner, F. & Massberg, S. Patrolling the vascular borders: platelets in immunity to infection and cancer. Nat Rev Immunol 19, 747–760, doi:10.1038/s41577-019-0202-z (2019).

42. Stark, K. & Massberg, S. Interplay between inflammation and thrombosis in cardiovascular pathology. Nat Rev Cardiol, doi:10.1038/s41569-021-00552-1 (2021).

43. Arunachalam, P. S. et al. Systems biological assessment of immunity to mild versus severe COVID-19 infection in humans. Science 369, 1210–1220, doi:10.1126/science.abc6261 (2020).

44. Tiedt, R., Schomber, T., Hao-Shen, H. & Skoda, R. C. Pf4-Cre transgenic mice allow the generation of lineage-restricted gene knockouts for studying megakaryocyte and platelet function in vivo. Blood 109, 1503–1506, doi:10.1182/blood-2006-04-020362 (2007).

45. Buch, T. et al. A Cre-inducible diphtheria toxin receptor mediates cell lineage ablation after toxin administration. Nat Methods 2, 419-426, doi:10.1038/nmeth762 (2005).

46. Muller, U. et al. Functional role of type I and type II interferons in antiviral defense. Science 264, 1918–1921, doi:10.1126/science.8009221 (1994).

47. Prigge, J. R. et al. Type I IFNs Act upon Hematopoietic Progenitors To Protect and Maintain Hematopoiesis during Pneumocystis Lung Infection in Mice. J Immunol 195, 5347–5357, doi:10.4049/jimmunol.1501553 (2015).

48. Swiecki, M., Gilfillan, S., Vermi, W., Wang, Y. & Colonna, M. Plasmacytoid dendritic cell ablation impacts early interferon responses and antiviral NK and CD8(+) T cell accrual. Immunity 33, 955–966, doi:10.1016/j.immuni.2010.11.020 (2010).

49. Yuan, L. et al. A role of stochastic phenotype switching in generating mosaic endothelial cell heterogeneity. Nat Commun 7, 10160, doi:10.1038/ncomms10160 (2016).

50. Mead, T. J. & Lefebvre, V. in Skeletal Development and Repair: Methods and Protocols (ed Matthew J. Hilton) 233–243 (Humana Press, 2014).

51. Love, M. I., Huber, W. & Anders, S. Moderated estimation of fold change and dispersion for RNA-seq data with DESeq2. Genome Biol 15, 550, doi:10.1186/s13059-014-0550-8 (2014).

52. Afgan, E. et al. The Galaxy platform for accessible, reproducible and collaborative biomedical analyses: 2016 update. Nucleic Acids Res 44, W3–W10, doi:10.1093/nar/gkw343 (2016).

53. Subramanian, A. et al. Gene set enrichment analysis: a knowledge-based approach for interpreting genome-wide expression profiles. Proc Natl Acad Sci U S A 102, 15545–15550, doi:10.1073/pnas.0506580102 (2005).

54. Mootha, V. K. et al. PGC-1α-responsive genes involved in oxidative phosphorylation are coordinately downregulated in human diabetes. Nature Genetics 34, 267–273, doi:10.1038/ng1180 (2003).

55. Metsalu, T. & Vilo, J. ClustVis: a web tool for visualizing clustering of multivariate data using Principal Component Analysis and heatmap. Nucleic Acids Res 43, W566-570, doi:10.1093/nar/gkv468 (2015).

